# Proteomic Profile of TGF-β1 treated Lung Fibroblasts identifies Novel Markers of Activated Fibroblasts in the Silica Exposed Rat Lung

**DOI:** 10.1101/431825

**Authors:** Mao Na, Xu Hong, Jin Fuyu, Xu Dingjie, Dominic Sales, Zhang Hui, Wei Zhongqiu, Li Shifeng, Gao Xuemin, Cai Wenchen, Li Dan, Zhang Guizhen, Zhang Bonan, Zhang Lijuan, Li Shumin, Zhu Ying, Wang Jin, Rui Mingwang, Ross Summer, Yang Fang

**Affiliations:** Basic Medical College, North China University of Science and Technology, Tangshan, China; Medical Research Center, North China University of Science and Technology, Tangshan, China; College of Traditional Chinese Medicine, North China University of Science and Technology, Tangshan, China; Center for Translational Medicine, Jane and Leonard Korman Respiratory Institute, Thomas Jefferson University, Philadelphia, PA, USA; School of public health, North China University of Science and Technology, Tangshan, China

**Keywords:** proteomics, fibroblast, silicosis, lung, pulmonary fibrosis

## Abstract

We performed liquid chromatography-tandem mass spectrometry (LC-MS/MS) on control and TGF-β1-exposed rat lung fibroblasts to identify proteins differentially expressed between cell populations. A total of 1648 proteins were found to be differentially expressed in response to TGF-β1 treatment and 196 proteins were expressed at ≥ 1.2 fold relative to control. Guided by these results, we next determined whether similar changes in protein expression were detectable in the rat lung after chronic exposure to silica dust. Of the five proteins selected for further analysis, we found that levels of all proteins were markedly increased in the silica-exposed rat lung, including the proteins for the very low density lipoprotein receptor (VLDLR) and the transmembrane (type I) heparin sulfate proteoglycan called syndecan 2 (SDC2). Because VLDLR and SDC2 have not, to our knowledge, been previously linked to the pathobiology of silicosis, we next examined whether knockdown of either gene altered responses to TGF-β1 in MRC-5 lung fibroblasts. Interestingly, we found knockdown of either VLDLR or SDC2 dramatically reduced collagen production to TGF-β1, suggesting that both proteins might play a novel role in myofibroblast biology and pathogenesis of silica-induced pulmonary fibrosis. In summary, our findings suggest that performing LC-MS/MS on TGF-β1 stimulated lung fibroblasts can uncover novel molecular targets of activated myofibroblasts in silica-exposed lung.

**Highlights:** We identified 196 proteins differentially expressed between control and TGF-β1 treated fibroblastsby LC-MS/MS.

Several proteins identified by LC-MS/MS were also found to be differentially expressed in whole lung tissues and isolated fibroblasts after chronic exposure to silica dust, including the very low density lipoprotein receptor (VLDLR) and the transmembrane type I heparan sulfate proteoglycan called syndecan 2

Knockdown of SDC2 or VLDLR markedly inhibited collagen production in MRC-5 fibroblasts, suggesting a novel pathogenic role for these proteins in myofibroblast biology.

## INTRODUCTION

Silicosis is a highly prevalent occupational condition that is caused by chronic inhalation of crystalline silica particulates (aerodynamic diameter < 5 μm) into the distal air spaces of the lung. Though this disease is largely preventable by implementing strict safety and health standards at work, silicosis remains one of the more common occupational disorders in many parts of the world, including China.

Although silica dust has been shown to injure many cell types in the lung it is generally believed that activated myofibroblasts play a central role in driving the development of disease. Central to this activation is signaling through the transforming growth factor β1 (TGF-β1) receptor, which leads to a cascade of events that trigger the transdifferentation of fibroblasts to myofibroblasts and induce the production of large quantities of collagen and other extracellular matrix materials (Gauldie *et al*., 1999; Harris *et al*., 2013; Jagirdar *et al*., 1996). Despite decades of work on TGF-β1 signaling in lung fibroblasts much remains to be learned about molecular mechanisms controlling myofibroblast growth and activation. Moreover, it is largely believed by many experts in the field that gaining additional insight into biological events downstream of TGF-β1 signaling could lead to the identification of new molecular targets for the treatment of silicosis and various other fibrotic lung conditions.

In our previous work, we utilized a 2-DE proteomic approach to identify novel biomarkers of silicosis (Xiaojun *et al*., 2016). However, it is now clear that using isobaric tags for relative quantification (iTRAQ) has several advances over the 2-DE approach including higher throughput quantification, decreased analytical time, and lower run-to-run variation (Rauniyar *et al*., 2014). With this in mind, the objective of this work was to utilize iTRAQ coupled liquid chromatography-tandem mass spectrometry (LC-MS/MS) to identify novel markers of activated rat lung fibroblasts and to utilize this information to assess whether similar markers were upregulated in the rat lung after chronic exposure to silica dust.

## MATERIALS AND METHODS

### Cell culture

Rat lung fibroblasts were isolated from 1–3 day old Wistar rats as previously described (Xu *et al*., 2012). Briefly, freshly isolated cells were culture for 3 to 4 passages and then transferred to low serum conditions (0.5% FBS) for 24 h. Following serum deprivation, cellwere treated with 5 ng/ml TGF-β1 (240-B; R&D Systems, Inc., Minneapolis, MN, USA) or vehicle controlfor 24 h and whole cell lysates were collected for later analysis (Xiaojun, et al., 2016). For each group, cells from 10 independent samples were pooled for proteomics analysis.

Fibroblasts used in our studies were derived from the lungs of control and silica-exposed rats(Seluanov *et al*., 2010) and the MRC-5 cell line. Fibroblasts were cultured in MEM containing 2% HEPES, 1% nonessential amino acid, and 10% FBS. Transient transfections were carried out using lipofectamine 2000 (Invitrogen, USA) according to the manufacturer’s recommendations. Knockdown was performed by exposing MRC-5 cells to 10 ng of siRNA targeting either very low-density lipoprotein receptor (VLDLR) or syndecan-2 (SDC2). Six hours after transfection, cells were washed and exposed to medium containing vehicle control or TGF-β1.

siRNAs used in our studies were derived from following sequences: 1) VLDL R-siRNA 1: GUGCAACAAUGGCCAGUGU dTdT; CACGUUGUUACCGGUCA CA dTdT; 2) VLDLR-siRNA 2: GCGAGUGCAUCCAUAAGAA dTdT; CGCU CACGUAGGUAUUCUU dTdT; 3) VLDLR-siRNA 3: GAUCGACAAUGUCUA UAAU dTdT; CUAGCUGUUACAGAUAUUA dTdT; 4) SDC2-siRNA 1: CCA CGACGCUGAAUAUACA dTdT; GGUGCUGCGACUUAUAUGU dTdT; 5) SD C2-siRNA 2: GUUGGUGUAUCGCAUGAGA dTdT; CAACCACAUAGCGUAC UCU dTdT; 6) SDC2-siRNA 3: GGAGAACGCAAACCAUCCA dTdT; CCUCU UGCGUUUGGUAGGU dTdT.

### Sample preparation for LC-MS/MS analysis

Proteins for LC-MS/MS were as prepared previously described (Wisniewski *et al*., 2009)(Promega, Fitchburg, Wisconsin, USA). Briefly, cell lysates were loaded onto a 10-kDa filter unit (Pall Corp., Port Washington, New York, USA), and filter was sequentially washed with UA (8 M urea in 0.1 M Tris-HCl;pH 8.5) and ABC (25 mMNH_4_HCO_3_) buffers. Proteins in collected samples were then denatured in 20 mM DTT at 50°C for 1 h and carboxyamidomethylated with 50 mM iodoacetamide (IAA) for 45 min. This step was followed by a digestion step using Trypsin Gold (1:50) at 37°C overnight. Peptide fragments were next desalted using Oasis HLB cartridges (Waters, Milford, Massachusetts, USA), dried by vacuum centrifugation (Thermo Fisher Scientific, Bremen, Germany) and again collected as a filtrate. Finally, samples were labelled with 4-plex iTRAQ reagent (AB Sciex) according to the manufacture’s protocol, with the control group labelled with tag 116 and TGF-β group with tag 117.

### High-performance liquid chromatography (HPLC) separation

Pooled mixtures of labelled samples were fractionated using a high-pH HPLC on a Waters Xbridge C18 column (4.6 mm × 250 mm, 3 μm). Samples were loaded onto a column equilibrated in buffer A1 (H2O; pH10), with an elution gradient of 5 – 25% buffer B1 (90% ACN; pH10) and a flow rate = 1 mL/min for 60 min. Flow-through was collected at 1minute intervals for a total of 60 fractions. Solutions were then dried, re-suspended in 0.1% formic acid, and pooled by concatenating fractions 1, 31; 2, 32; and so on. Analyses were performed on just odd number fractions.

### LC-MS/MSanalysis

Each fraction was analysed with a reverse-phase-C18 self-packed capillary LC column (75 μm × 100 mm), with an elution gradient of 5% – 30% buffer B2 (0.1% formic acid, 99.9% ACN; flow rate = 0.3 μL/min) for 40 min. A Triple TOF 5600 mass spectrometer was used to analyze the eluted peptides, with each fraction run in duplicate. The MS data were acquired using the high-sensitivity mode with the following parameters: 30 data-dependent MS/MS scans per full scan; full scans acquired at a resolution of 40,000 and MS/MS scans at a resolution of 20,000; rolling collision energy, charge state screening (including precursors with a charge state of +2 to +4), and dynamic exclusion (exclusion duration 15 s); and a MS/MS scan range of 100 – 1800 m/z, with a scan time of 100 ms.

### Date analysis

Data acquired from LC-MS/MS were analysed using the Mascot software (version 2.4.1, Matrix Science, London, UK). Proteins were identified by searching peptide spectral matches against the Swissprot_2014_07 database (taxonomy: Rattus, containing 7,906 sequences). The parameters utilized included trypsin as the digestion enzyme, two or fewer missed cleavage sites and cysteine carbamidomethylation (57.02146) as a fixed modification. The precursor ion mass tolerance and the fragment ion mass tolerance were set to ppm and 0.05 Da respectively. Protein identifications from Mascot were validated using the Scaffold Proteome Software (version 4.3.3, Proteome Software Inc., Portland, OR). Peptide identifications were accepted if false discovery rate (FDR) was less than 1.0% and if at least 2 unique peptides were identified from the same protein. iTRAQ, Scaffold Q+ was used for Label Based Quantification (TMT, iTRAQ, SILAC, etc.) of peptides and proteins. For each channel, the acquired reporter ion intensities were normalized by the sum of all reporter ion intensities of that channel. Normalized intensities were then used to calculate the relative protein abundance and quantify protein ratios(Nesvizhskii *et al*., 2003). Proteins identified from these analyses were further analysed to determine function, ontology and location using the PANTHER classification system (Protein Analysis Through Evolutionary Relationships; http://www.pantherdb.org/) (Mi *et al*., 2013).

### Silicosis model

Male Wistar rats were purchased from Vital River Laboratory Animal Technology Co. Ltd. (SCXY 2009-0004; Beijing, China) and all experiments were performed in accordance with the regulations set by the Committee on the Ethics of North China University of Science and Technology. Silica dust was delivered to rats using a HOPE MED 8050 exposure control apparatus (HOPE Industry and Trade Co. Ltd, Tianjin, China) with the SiO2(s5631, Sigma-Aldrich, St. Louis, MO, USA) concentration maintained at 2,000 mg/m^3^ (Liu *et al*., 2017). At experimental endpoints, bronchoalveolar lavage was performed with 0.9% saline and whole lung tissues were then snap-frozen in liquid nitrogen.

### Immunohistochemistry (IHC)

Paraffin-embedded tissue sections were used for IHC. Endogenous peroxidases were quenched with 3% H_2_O_2_ and antigen retrieval was performed using a high-pressure method with deparaffinised sections. The samples were then incubated with primary antibodies against collagen V(COL V, A1515, ABclonal Biotech, Wuhan, China), collagen XI (COL XI, DF3553, Affinity Biosciences, Cincinnati, OH, USA), VLDLR (TA309928, OriGene Technologies, Rockville, MD, USA) or SDC2 (A1810, ABclonal Biotech, Wuhan, China) at 4°C overnight. The following morning, tissue sections were incubated with secondary antibody conjugated with the horseradish peroxidase enzyme (PV-6000; Beijing ZhongshanJinqiao Biotechnology Co. Ltd, China) at 37°C for 20 min. Immunoreactivity was visualised with using the DAB substrate (ZLI-9018; ZSGB-BIO, Beijing, China) and the nuclei were stained with hematoxylin.

### Western blot analysis

Total protein levels were quantified using a Bradford assay (PC0020; Solarbio, China) as previously described. Protein lysates (20 μg/lane) were separated on a 13% gel by SDS-PAGE and were electro-transferred onto polyvinylidene fluoride (PVDF) membranes (Amervehicle Biosciences). The membranes were blocked with 5% non-fat milk and incubated with primary antibodies against COL V, COL XI, Vascular cell adhesion protein (VCAM, ab134047, Abcam, Cambridge, MA, USA), Transmembrane protein 214 (TM214, TS306822, OriGene Technologies, Rockville, MD, USA), VLDLR, SDC-2 or α-SMA at 4°C overnight. Membranes were next washed in TBST and incubated with peroxidase-labelled affinity-purified anti-rabbit/mouse IgG (H + L) secondary antibody(074–1506/074–1806;Kirkegard and Perry Laboratories). Protein bands were visualised using ECL™ Prime Western Blotting Detection Reagent (RPN2232; GE Healthcare, Hong Kong, China) and expressed as a fold change relative to α-Tubulin (Tub α, AF7010; Affinity Biosciences, Cincinnati, OH, USA).

### Statistical analysis

Comparisons between two groups were performed using independent sample *t*-test. Comparisons between multiple independent groups were done using a one-way ANOVA followed by a post-hoc analysis with the Bonferroni test. Values were expressed as a means ± Std, with p-values less than 0.05 considered as statistically significant.

## RESULTS

### Proteomic profile differs between control and TGF-β1 lung fibroblasts

To identify proteins differentially expressed between control and activated fibroblasts, we performed iTRAQ coupled with LC-MS/MS on rat lung fibroblasts treated with or without TGF-β1 for 24 h. Using scaffold integration, we observed a total of 1648 proteins differentially expressed between control and TGF-β1 treated cells. Moreover, 196 of these proteins were expressed at ≥ 1.2 fold-change (Tables S1 and S2) and 20 proteins exhibited a ≥ 1.5 fold-change (Table 1) relative to control.

**Table 1.** Subset of differentially expressed proteins in cultured TGF-β-induced myofibroblasts relative to the untreated control.

| **No.** | **Protein name** | **Accession Number** | **Molecular Weight** | **Fold change** |
| --- | --- | --- | --- | --- |
| 1 | Collagen alpha-1(II) chain | CO2A1_RAT | 135 kDa | 1.60 |
| 2 | Cluster of Histone H2A type 1 | H2A1_RAT [4] | 14 kDa | 1.70 |
| 3 | Collagen alpha-1(V) chain | CO5A1_RAT | 184 kDa | 1.50 |
| 4 | Protein S100-A11 | S10AB_RAT | 11 kDa | 1.59 |
| 5 | Vascular cell adhesion protein 1 | VCAM1_RAT | 81 kDa | 1.55 |
| 6 | Collagen alpha-1(XI) chain | COBA1_RAT | 181 kDa | 1.70 |
| 7 | Cluster of Thymosin beta-4 | TYB4_RAT | 5 kDa | 2.50 |
| 8 | Cluster of Haemoglobin subunit beta-1 | HBB1_RAT [2] | 16 kDa | 2.25 |
| 9 | Haemoglobin subunit alpha-1/2 | HBA_RAT | 15 kDa | 1.90 |
| 10 | Oxidised low-density lipoprotein receptor 1 | OLR1_RAT | 42 kDa | 1.55 |
| 11 | Transmembrane protein 214 | TM214_RAT | 77 kDa | 1.50 |
| 12 | Very low-density lipoprotein receptor | VLDLR_RAT | 97 kDa | 1.55 |
| 13 | Syndecan-2 | SDC2_RAT | 22 kDa | 1.90 |
| 14 | Serum albumin | ALBU_RAT | 69 kDa | 0.59 |
| 15 | Histone H4 | H4_RAT | 11 kDa | 0.25 |
| 16 | Corticosteroid 11-beta-dehydrogenase isozyme 1 | DHI1_RAT | 32 kDa | 0.65 |
| 17 | Collectin-12 | COL12_RAT | 82 kDa | 0.60 |
| 18 | Cluster of Neurotrimin | NTRI_RAT [2] | 38 kDa | 0.65 |
| 19 | Programmed cell death protein 4 | PDCD4_RAT | 52 kDa | 0.55 |
| 20 | Plasminogen receptor (KT) | PLRKT_RAT | 17 kDa | 0.60 |

### Bioinformatic analyses of differentially expressed proteins in control and TGF-β1 stimulated lung fibroblasts

Proteins whose expression changed most significantly after exposure to TGF-β1 were classified into functional classes using the PANTHER analysis. Gene ontology (GO) analysis classified proteins into three distinct categories entitled: 1) molecular function; 2) biological process; and 3) components of the cell. Among these categories most proteins were belonged to molecular functions group (Table S3), and the major subgroups within this category included: 1) catalytic activity (40%); 2) protein binding (26%), and 3) structural activity (20%). In the biological processes category (Table S4), most proteins belonged to the subgroups: 1) cellular process (26%); 2) metabolic process (22%), or 3) cellular component/biogenesis (10%) subgroup; and proteins within the components of a cell group belonged to 1) cell part (40%); 2) cellular organelle (25%); and 3) macromolecular complexes (18%) (Table S5).

Within PANTHER, class ontology was also performed (Table S6), with the largest number of proteins assigned to the following categories:1) nucleic acid binding (18%); 2) hydrolase (10%); 3) enzyme modulator and transferase (each 7%); 4) cytoskeletal protein (6%); 5) signalling molecule (6%); and 7) transfer/carrier protein (6%). Pathway analysis showed that all proteins that were differentially expressed in response to TGF-β1fell into one of the following signalling pathways: 1) integrin; 2) angiogenesis; 3) CCKR; 4)conadotropin-releasing hormone receptor; 5) p53; 6) Alzheimer disease-presenilin; 7) blood coagulation; 8) cadherin; 9) cholesterol biosynthesis; 10) FGF or 11) Wntsignalling pathways (Table S7).

### Proteinsincreased in TGF-β1 stimulated lung fibroblasts were also increased in the silica-exposed rat lung

To determine whether in vitro findings were representative of changes in vivo, we next performed IHC and western blot analysis on control and silica-exposed lung tissues for those proteins whose expression changed most significantly (≥1.5 fold increase) in response to TGF-β1 treatment. However, our analysis was limited to proteins in which a commercially available antibody for IHC or WB was available; this included the proteins COLV, COL XI, VCAM1, TM214, VLDLR, and SDC2 for protein identification.

As shown in Figure 1A, van Gieson (VG) staining confirmed the ability of our model to induce severe fibrotic responses in the rat lung as demonstrated by an increase in extracellular matrix deposition and the large number of silicotic nodules. Western blot analysis also showed an increase in COLI and α-SMA in whole lung tissues and primary lung fibroblasts after silica exposure (Figure 1B).

**Figure 1.**
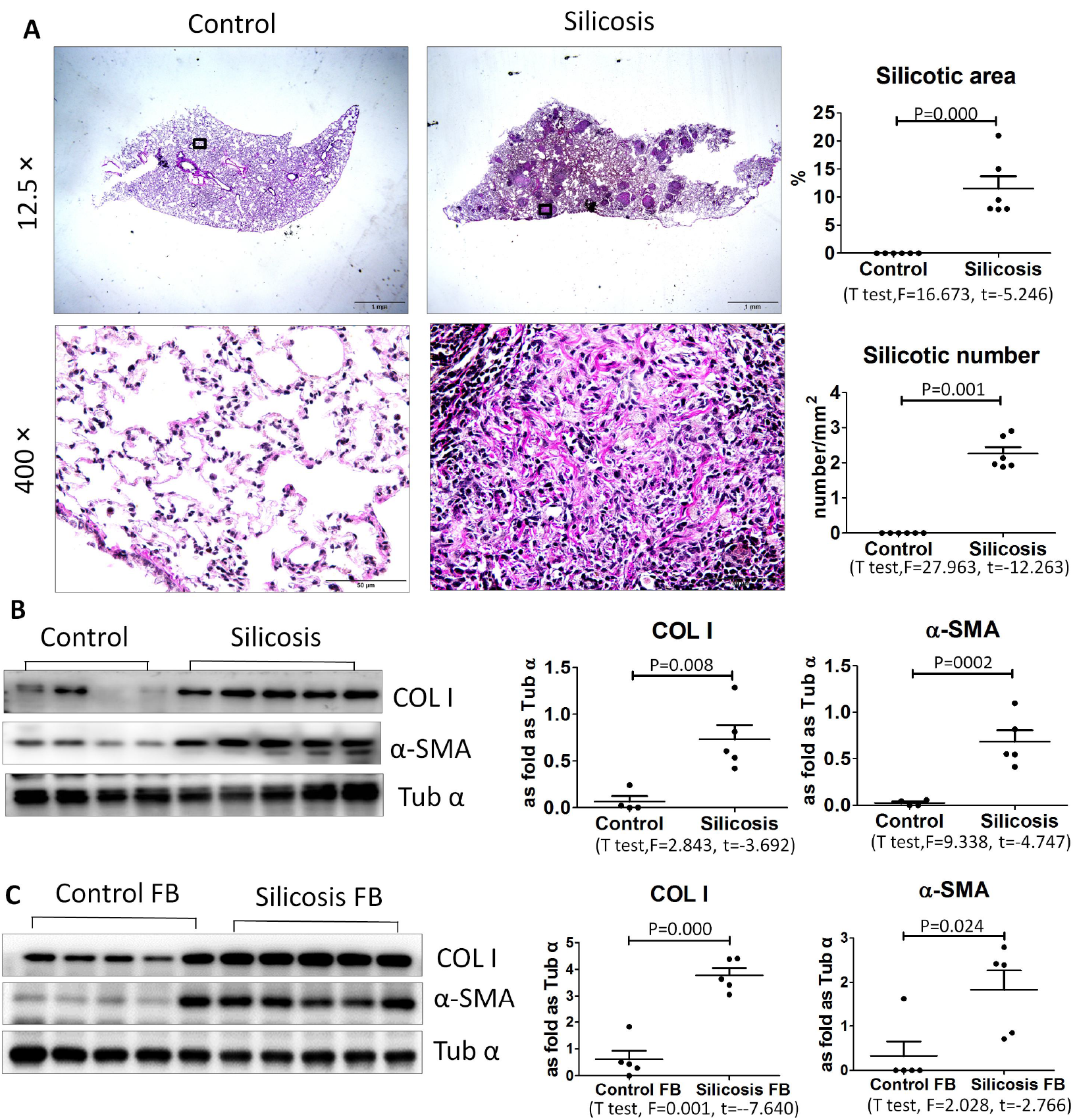
Histological examination of control and silica-exposed rat lung. A. Low (top) and high (bottom) power images of control and silica-exposed lung after van Gieson (VG) staining. Right) Quantitative assessment of area and number silicotic nodules in the lungs of control and silica-exposed rats. Western blot analysis for collagen I (COLI) and a-smooth muscle actin (α-SMA) in whole lung tissues and isolated fibroblasts from control and silica-exposed ratsAbbreviations: C, control; S, silicosis. Statistical significance was assessed using the Student’s t-test.

As shown in Figure 2, Western blot analysis revealed a marked increase in the expression of COL V, COL XI, and VCAM1 in the lungs of silica exposed rats, including whole lung tissues and freshly isolated lung fibroblasts. Moreover, IHC staining of the lung showed the elevated levels of COL V and COL XI was confined mostly to the interstitium of the lung, particularly in areas with evidence of active inflammation and fibrosis. Despite multiple attempts, staining for VCAM1 was detected in either the control or the silica-exposed rat lung, indicating that our antibody could not be used for IHC.

**Figure 2.**
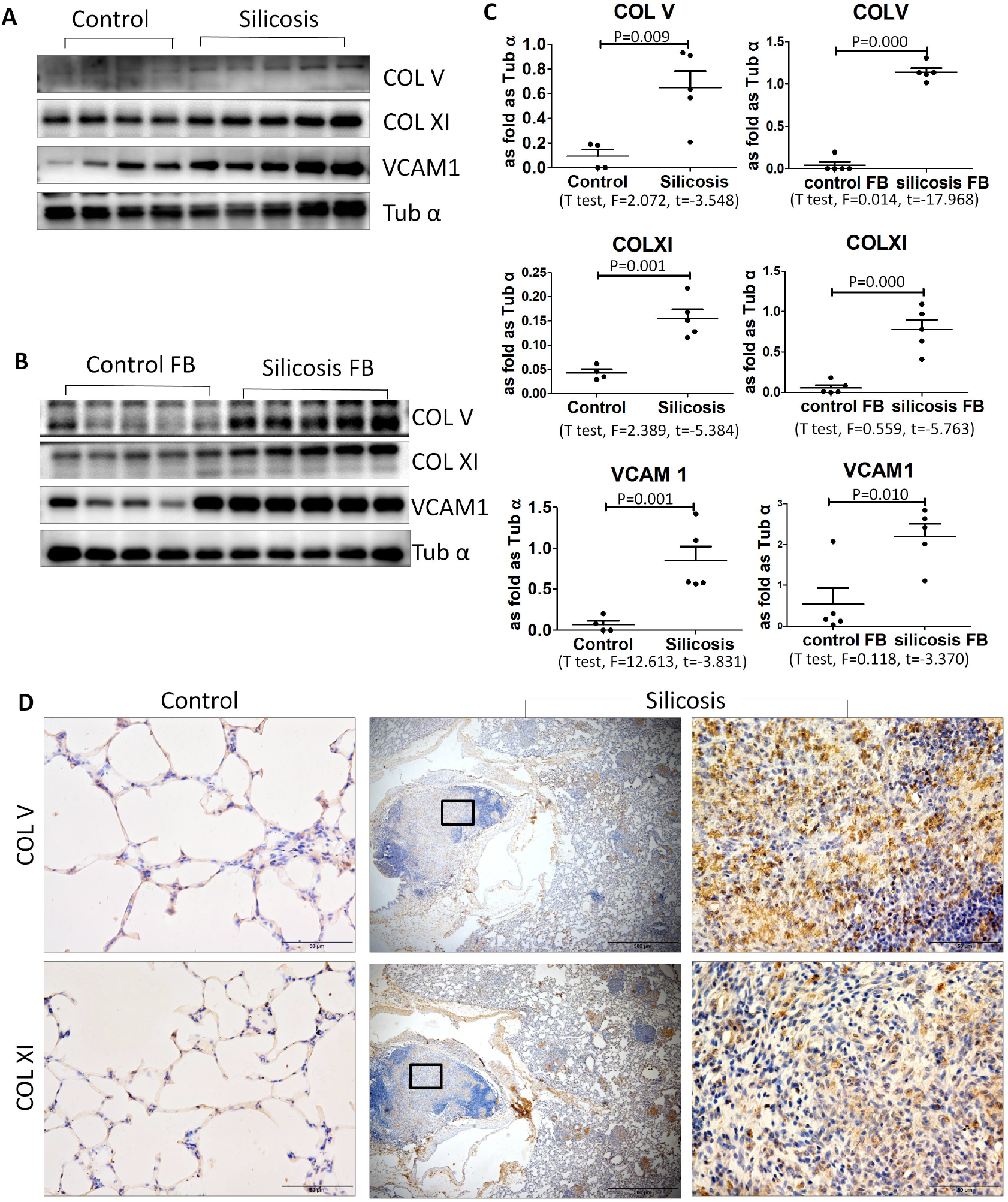
COL V, COLXI and VCAM1levels are increased in the rat lung after silicaexposure. A-C) Western blot analysis for COL V, COLXI and VCAM1 in control and silica exposed rat lung tissues. Quantitative analyses of immunoblots are shown in (C). D) IHC for expression COL V and COLXI in lung tissues from control (left) andsilica-exposed rats. Statistical significance was assessed using the Student’s t-test.

In addition to the above proteins, we found that levels of VLDLR and SDC2 were also markedly increased by western blot in whole lung tissues and isolated fibroblasts after silica exposure. (Figure 3). Significant increases in VLDLR and SDC2 expression were also observed by IHC in the silica-exposed lung.

**Figure 3.**
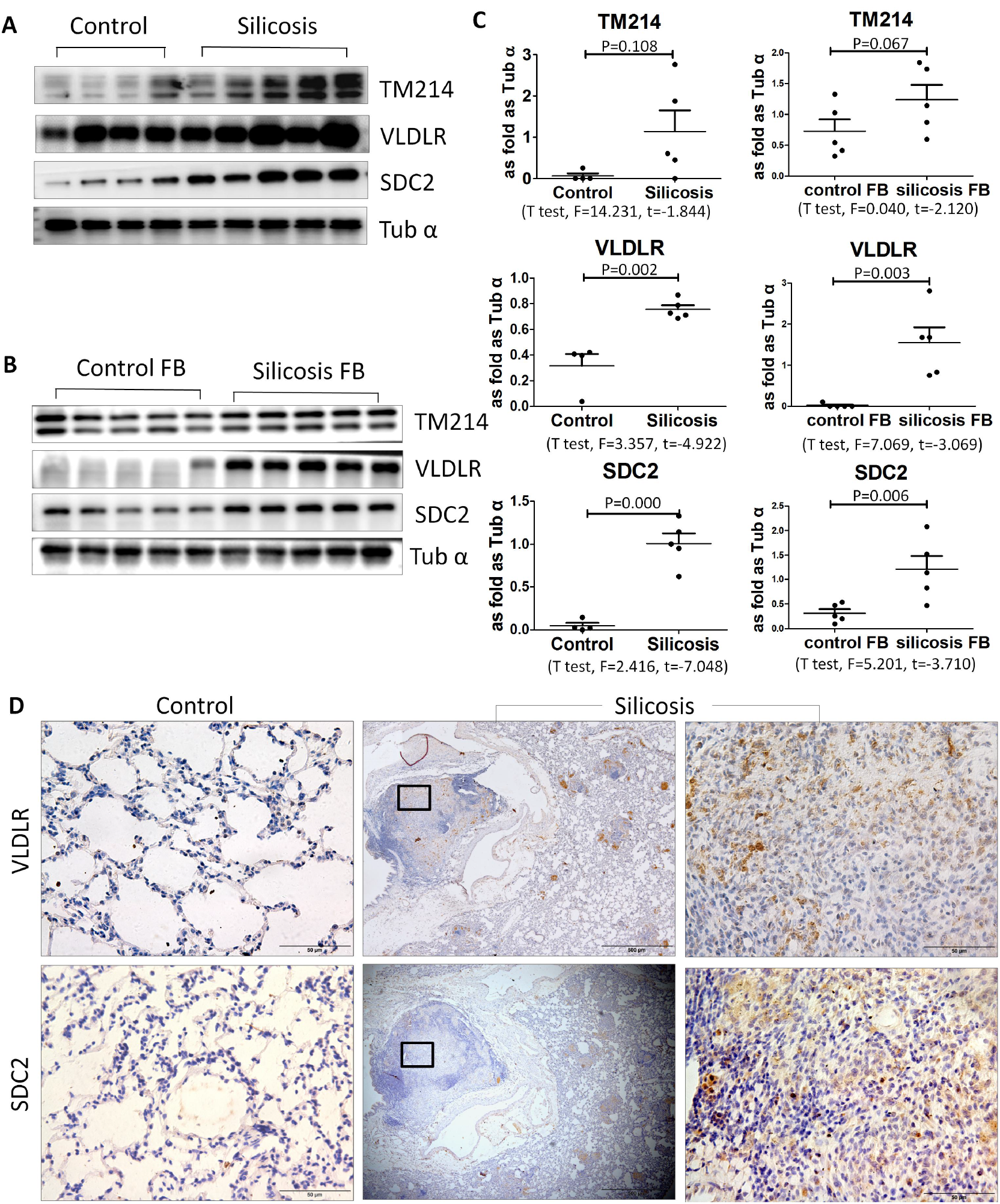
TM214, VLDLR and SDC2 expression is increased in the rat lung after silica exposure. A-C) Western blot analysis for TM214, VLDLR and SDC2 in control and silica exposed rat lung tissues. Quantitative analyses of immunoblots are shown in (C).D) IHC for expression VLDLR and SDC2 in lung tissues from control (left) andsilica-exposed rats. Statistical significance was assessed using the Student’s t-test.

### Knockdown of VLDLR or SDC2 reduces collagen deposition in TGF-β1stimulated fibroblasts

Although not entirely surprising that collagen levels COL V and COL XL) and VCAM expression(Agassandian *et al*., 2015) were increased in fibrotic lung tissues we were intrigued by the observation that levels of VLDLR and SDC2 were also markedly elevated in the lung after silica exposure. Even after an exhaustive search, we were unable to find any reports linking either VLDLR or SDC2 to silicosis in any tissues. Based on this, we speculated whether changes in the expression of VLDLR or SDC2 can influence fibrotic responses to TGFβ1 in lung fibroblasts. To test this, we next performed siRNA knockdown of VLDLR or SDC2 in MRC-5 cells using several different siRNA probes. Because the quantity of gene knockdown varied with different siRNAs we utilized only siRNAs demonstrating the most effective knockdown in our studies; this was siRNA 2 and 3 for VLDLR, and siRNA 3 for SDC2 (Figure 4). As shown in Figure 4, we found that knockdown of VLDLR markedly reduced COL I and α-SMA expression in TGFβ1 treated MRC-5 fibroblasts. Moreover, we also found that knockdown of SDC2 significantly suppressed COLI expression, although non-significant decreases in α-SMA levels were seen in TGFβ1 treated cells.

**Figure 4.**
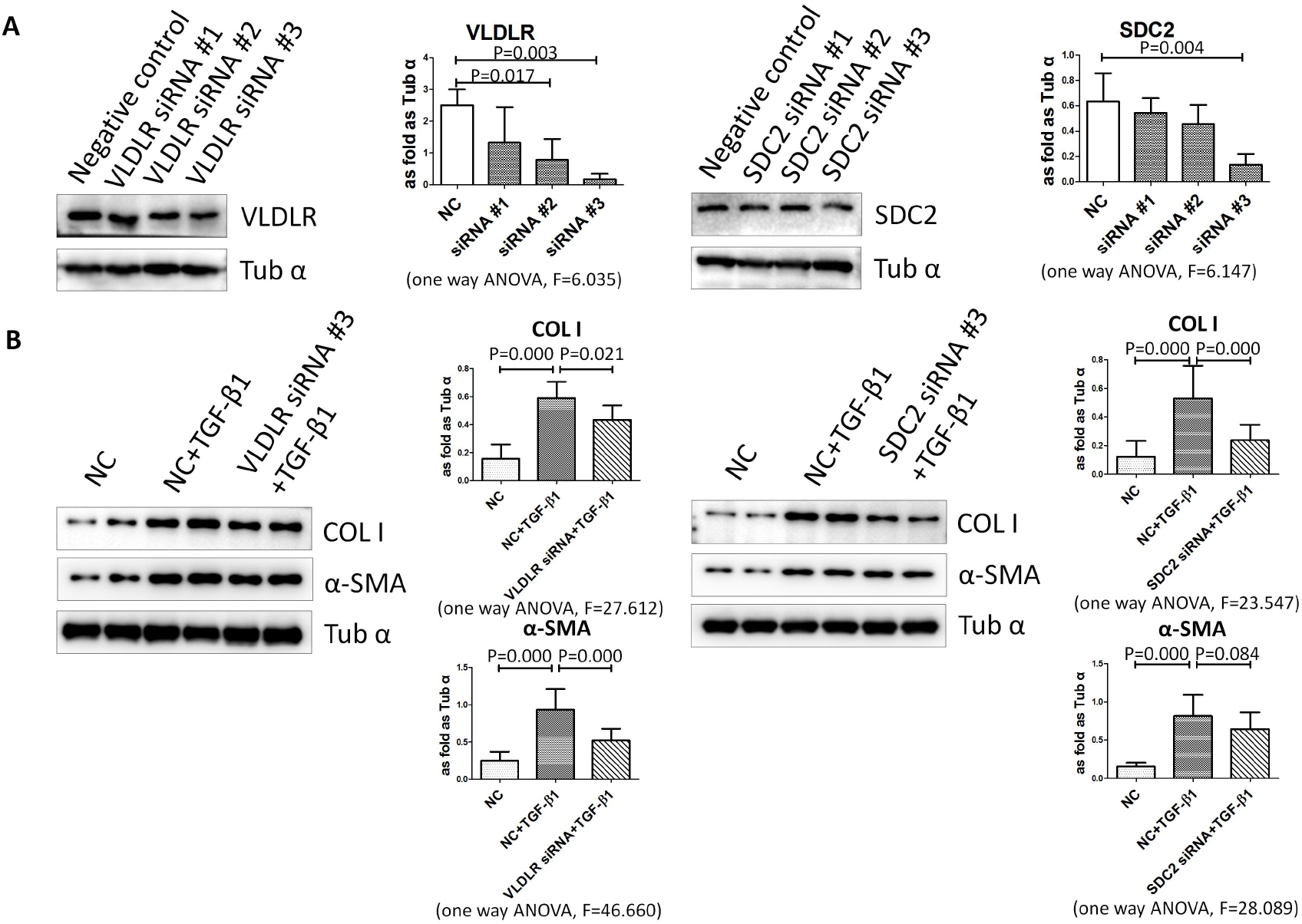
siRNA knockdown or VLDLR or SDC2 in lung fibroblasts reduces TGF-β1-induced collagen. A) Effectiveness of different siRNA probes in knockdown expression of VLDLR or SDC2. B) siRNA knockdown of VLDLR reduces expression of COL I and α-SMA in TGF-β1 treated MRC-5 fibroblasts. siRNA knockdown of SDC2 reduces expression of COL1 and causes a non-significant reduction in α-SMA in TGF-β1 treated MRC-5 fibroblasts.Statistical significance was assessed using the one way ANOVA.

## DISCUSSION

Biomarker discovery can be accomplished using a variety of proteomic approaches and biological specimens, including urine, serum or lysates from whole organ digests. However, for research in solid organs, such as the heart, brain, kidney or lung, it is often difficult to obtain sufficient quantities of patient samples for proteomic work. Moreover, tissue lysates, are composed of a conglomeration of different cell types, such as epithelial cells, smooth muscle cells, fibroblasts, immune cells and endothelialium, making it difficult to uncover novel biological mechanisms in individual cell types. Thus, *in vitro* models of individual cell populations are often employed to identify novel candidate biomarkers of functional processes *in vivo* (Paul *et al*., 2013).

In this study, we performed iTRAQ-coupled LC-MS/MS analysis on rat lung fibroblasts treated with or without TGF-β1 to identify novel markers of activated myofibroblasts in the silica-exposed lung. From these studies, we identified many proteins differentially expressed and 196 proteins expressed at ≥ 1.2 fold relative to control. Importantly, many of these proteins were novel markers and had not previously been linked to TGF-β1 signalling in the lung, including several whose functional classes have been intimately tied to myofibroblast activation such as integrin and angiogenesis pathways as well as those less clearly associated with myofibroblast biology such as nucleic acid binding, hydrolase activity and transferase activity.

An important finding in this study was the observation that some of the proteins we identified in our proteomics analyses were also differentially expressed in the lung after silica exposure. For example, we found that levels of COL XI and COL V were markedly increased in the silicotic lung, although these findings were not surprising given the fact that collagen production is a well-recognized by product of TGF-β1 signalling(Raglow *et al*., 2015). However, we also found that VCAM were significantly increased in the lung after silica exposure. The former findings is also not unexpected given that transcript and protein levels of VCAM are reported to be increased in fibroblastic foci of IPF lungs (Agassandian, et al., 2015).

Two other proteins found to be differentially expressed in our proteomic analyses were VLDLR and SDC2. We were intrigued by these observations because our review of the literature had not previously associated these proteins with silicosis. Western blot analysis and IHC confirmed that both proteins were increased in whole lung tissues and isolated fibroblasts from the lungs of silica-exposed rats. Interestingly, VLDLR expression is known to be increased in the fibrotic caps of atherosclerotic lesions in systemic blood vessels, suggesting it may be involved in more than just lipid trafficking (Eck *et al*., 2005). In IPF patient lung tissues, SDC2 was found to be over-expressed in two reports (Chen *et al*., 2004; Ruiz *et al*., 2012). More importantly, we found that knocking down the expression of either VLDLR or SDC2 effectively reduced collagen production, supporting the notion that these proteins play a role in the pathobiology of pulmonary fibrosis.

In conclusion, iTRAQ coupled LC-MS/MS of TGF-β1-induced fibroblasts can be utilized to identify novel markers of silica-induced lung fibrosis. Future mechanistic studies will be needed to uncover whether individual proteins are simply a marker of disease or play a biological role in the onset or progression of pulmonary fibrosis.

## Ethics approval

All animal experiments were reviewed and approved by the Institutional Animal Care and Use Committee at the North China University of Science and Technology University prior to the initiation of any studies.

## Consent for publication

Not applicable.

## Authors’ contributions

FY and HX designed the study. MN, JF, ZH, XD, YY, MN, GX, and GY carried out the experimental work, analyzed the data, and drafted the manuscript. RS participated in the design of the study and critically reviewed the manuscript and provided intellectual input. RS and RM helped write and critically reviewed the manuscript and provided intellectual input. WZ, ZB, LS, LS and WJ conceived the study, and participated in coordination, and helped in drafting the manuscript. All authors read and approved the final manuscript.

## Acknowledgements

Not applicable.

---

**Table S1.**
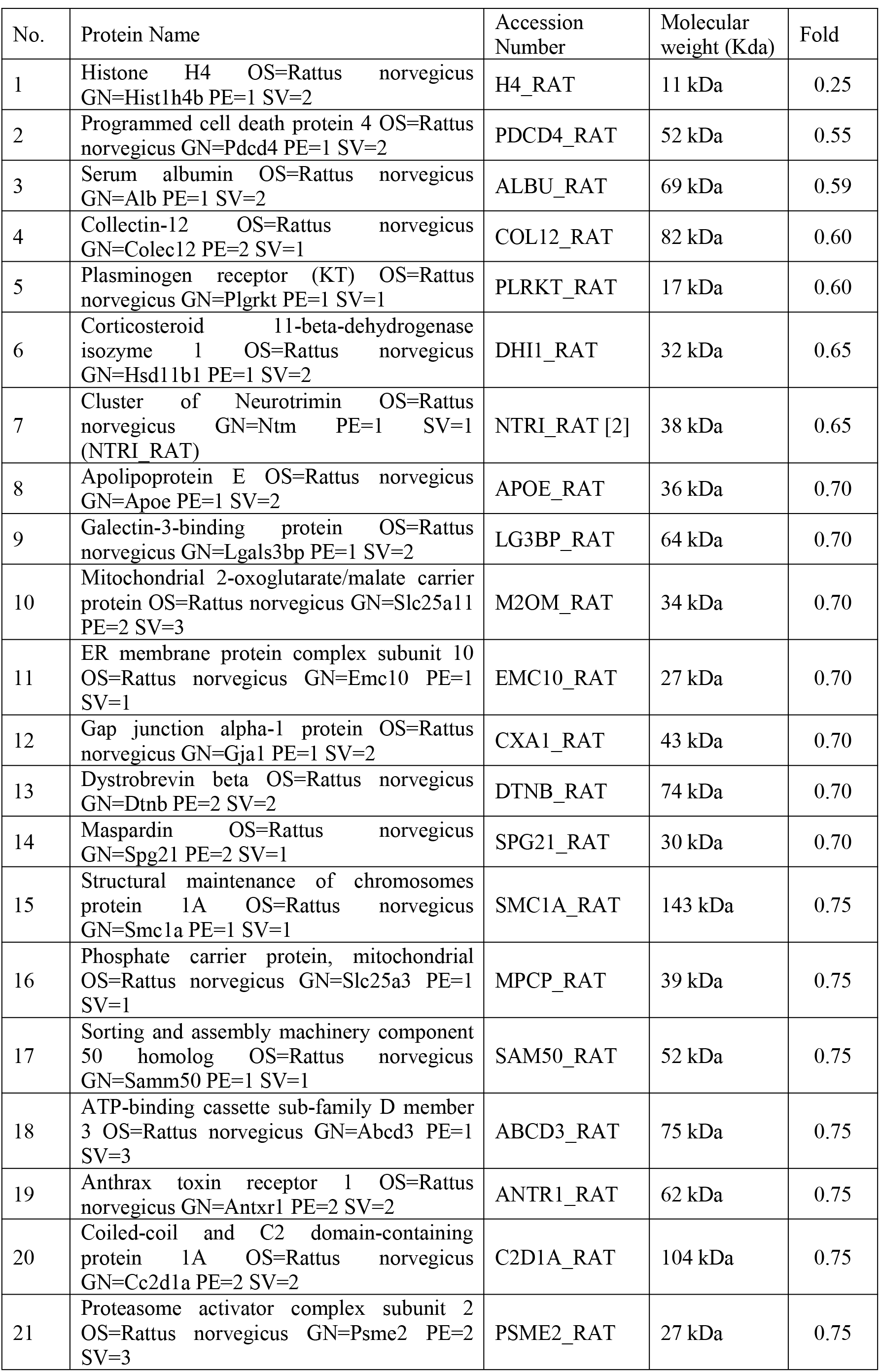
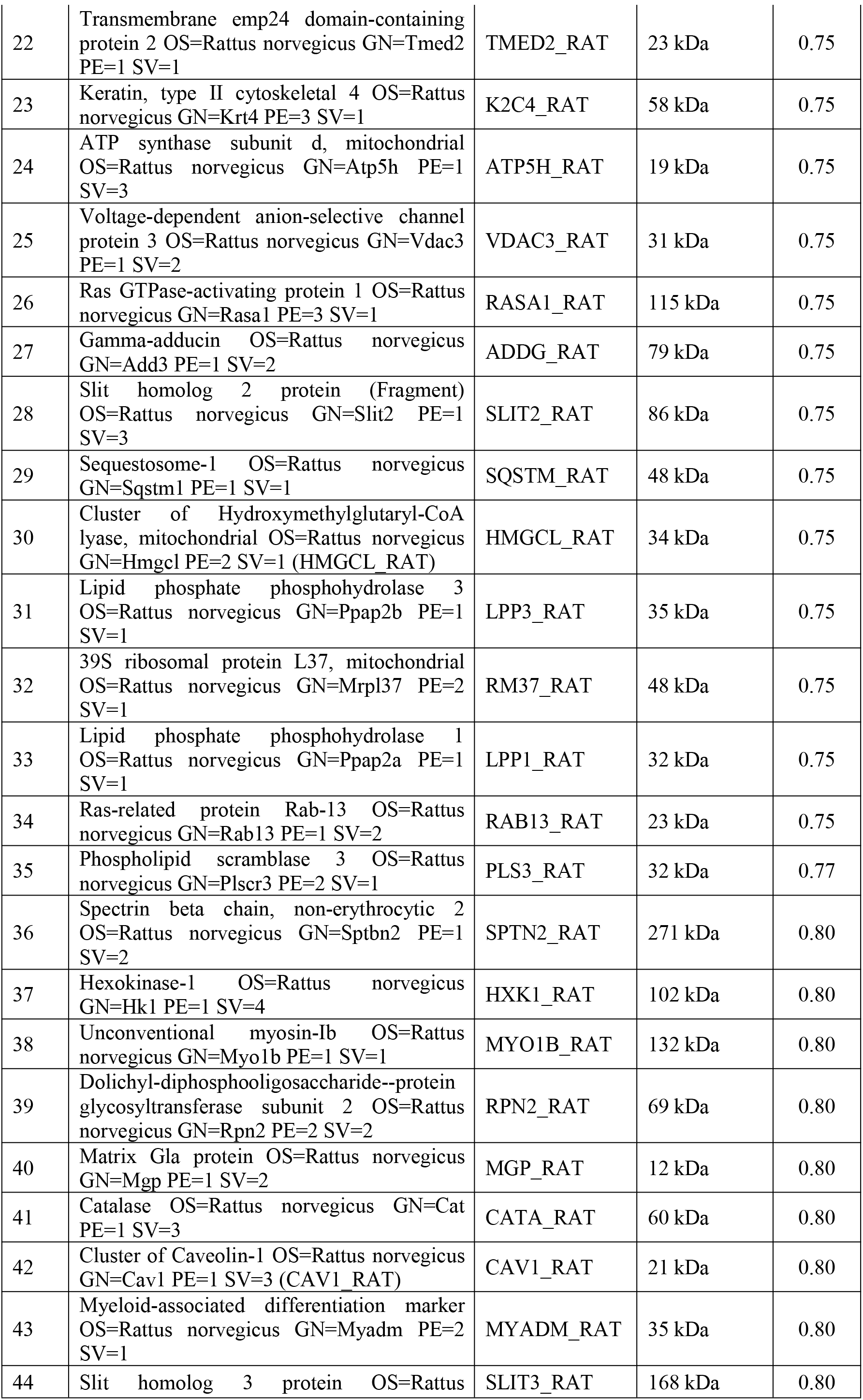
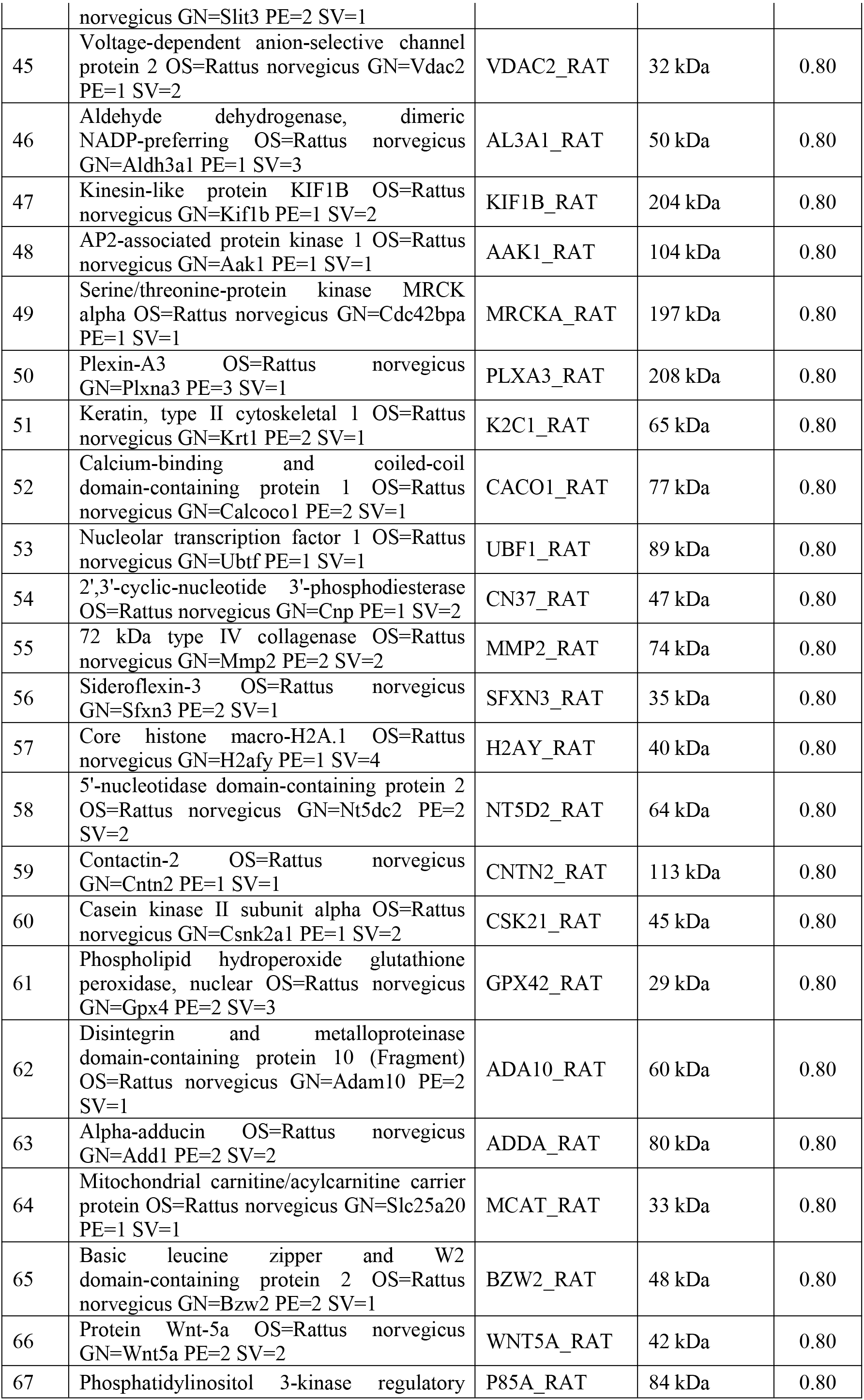
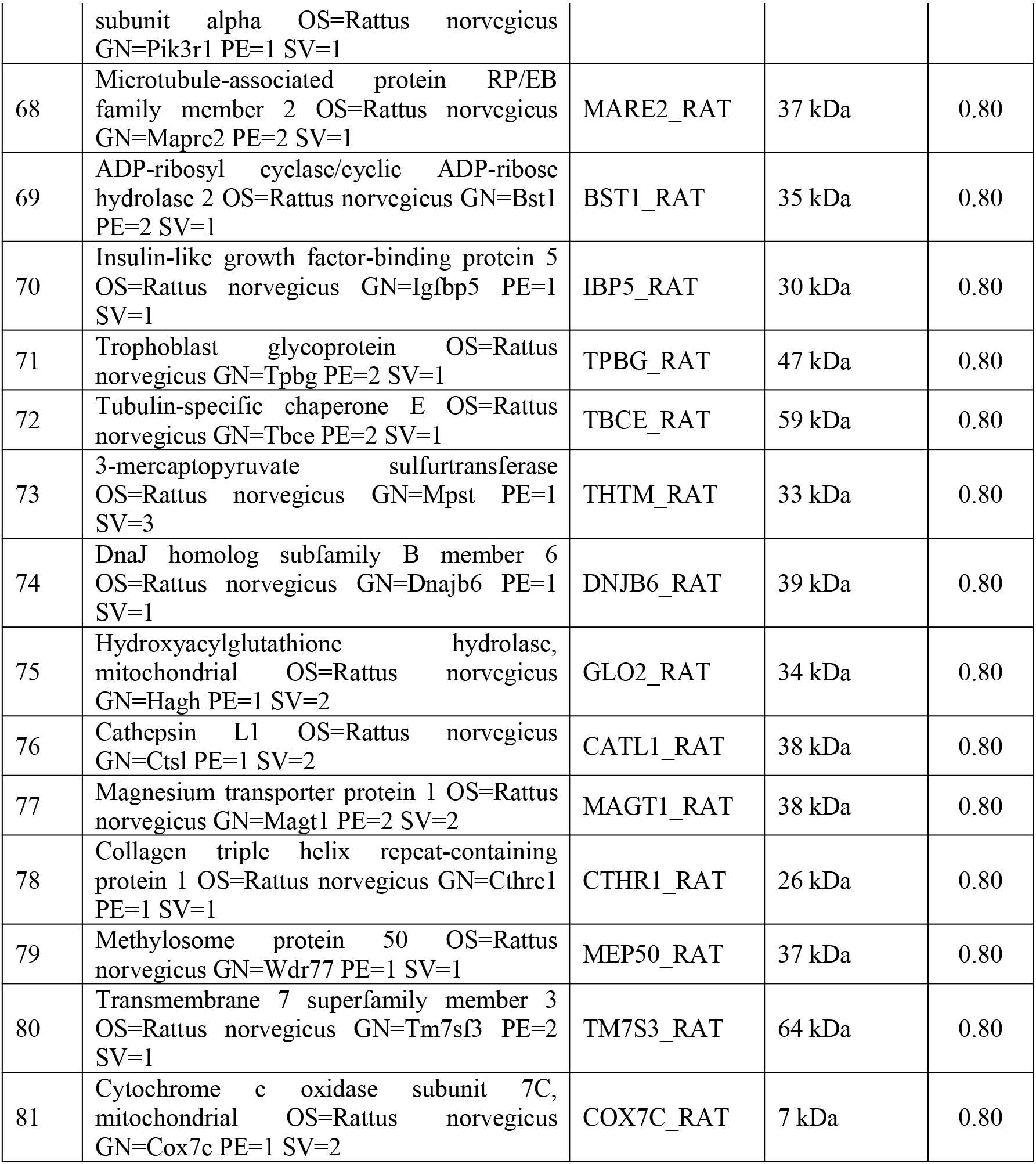
The down-regulation of differential proteins in fibroblasts induced by TGF-β1.

**Table S2.**
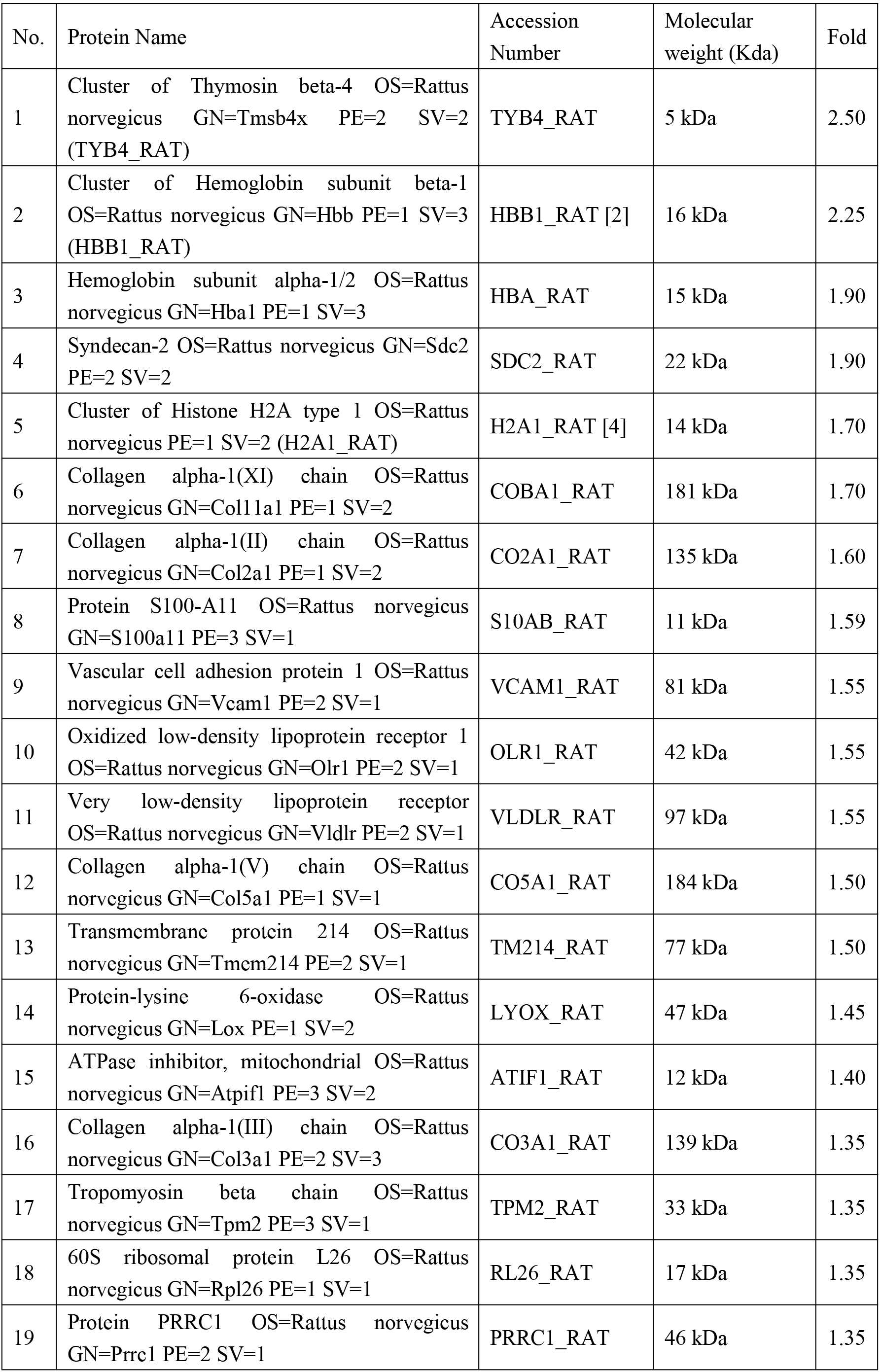
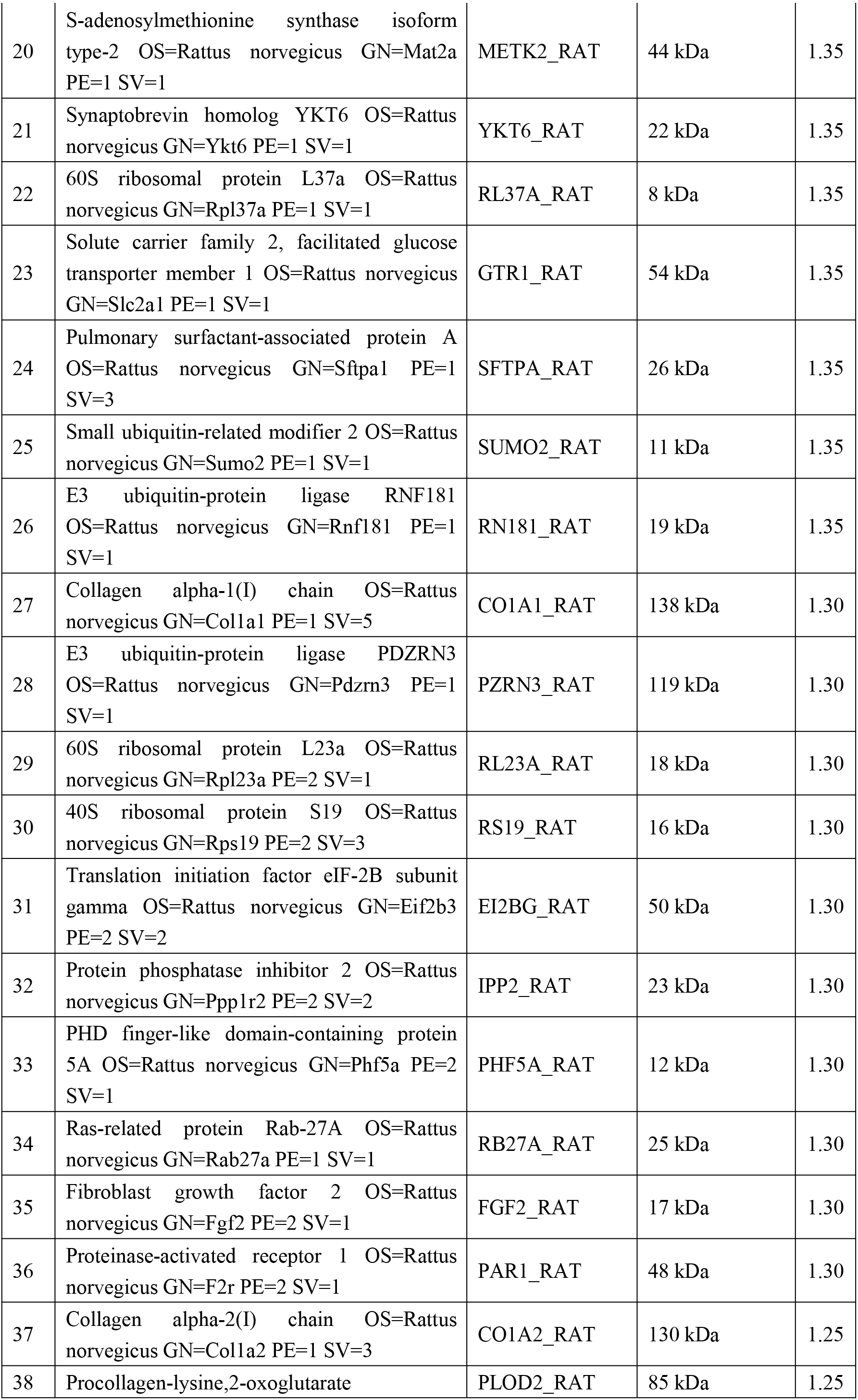
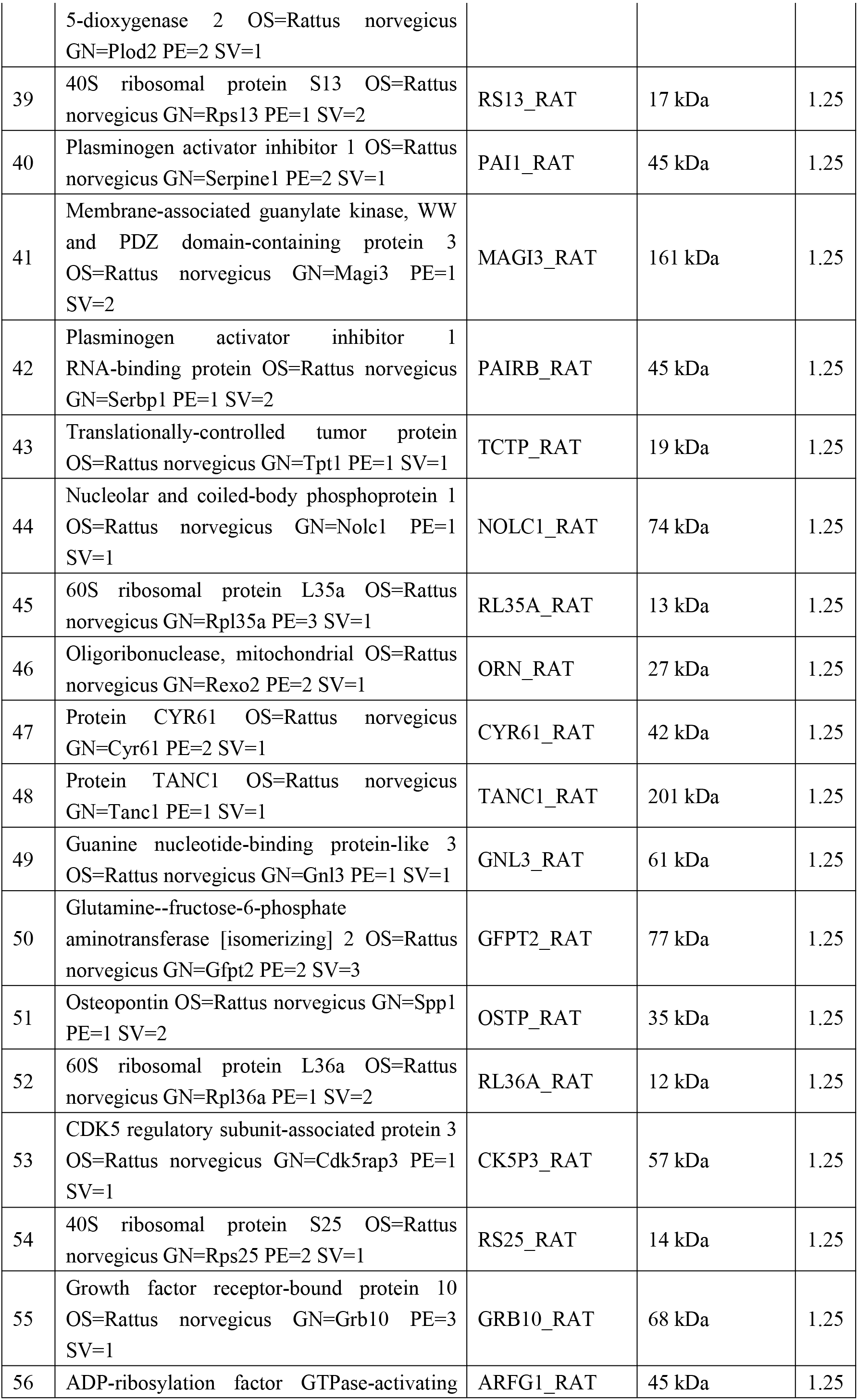
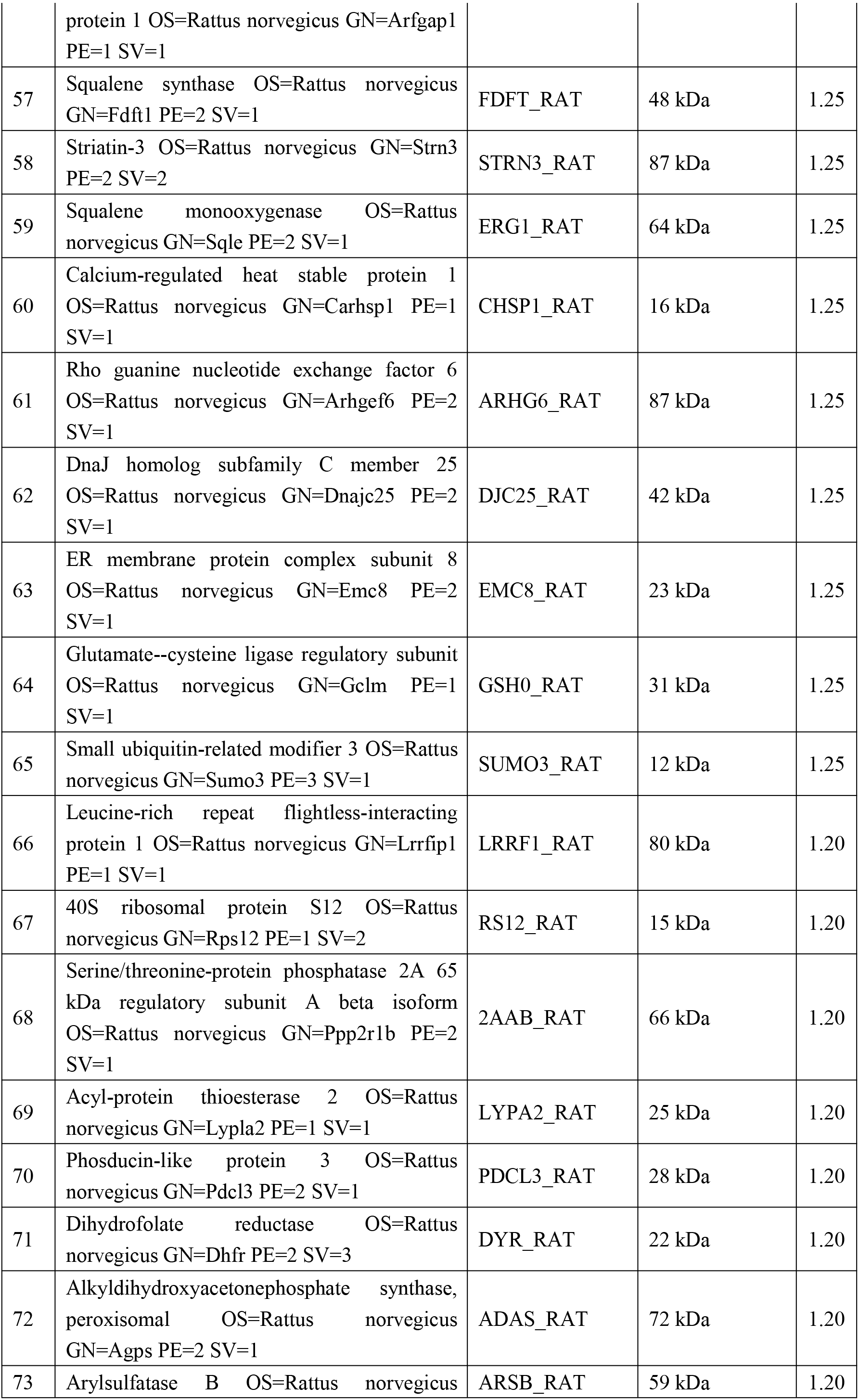
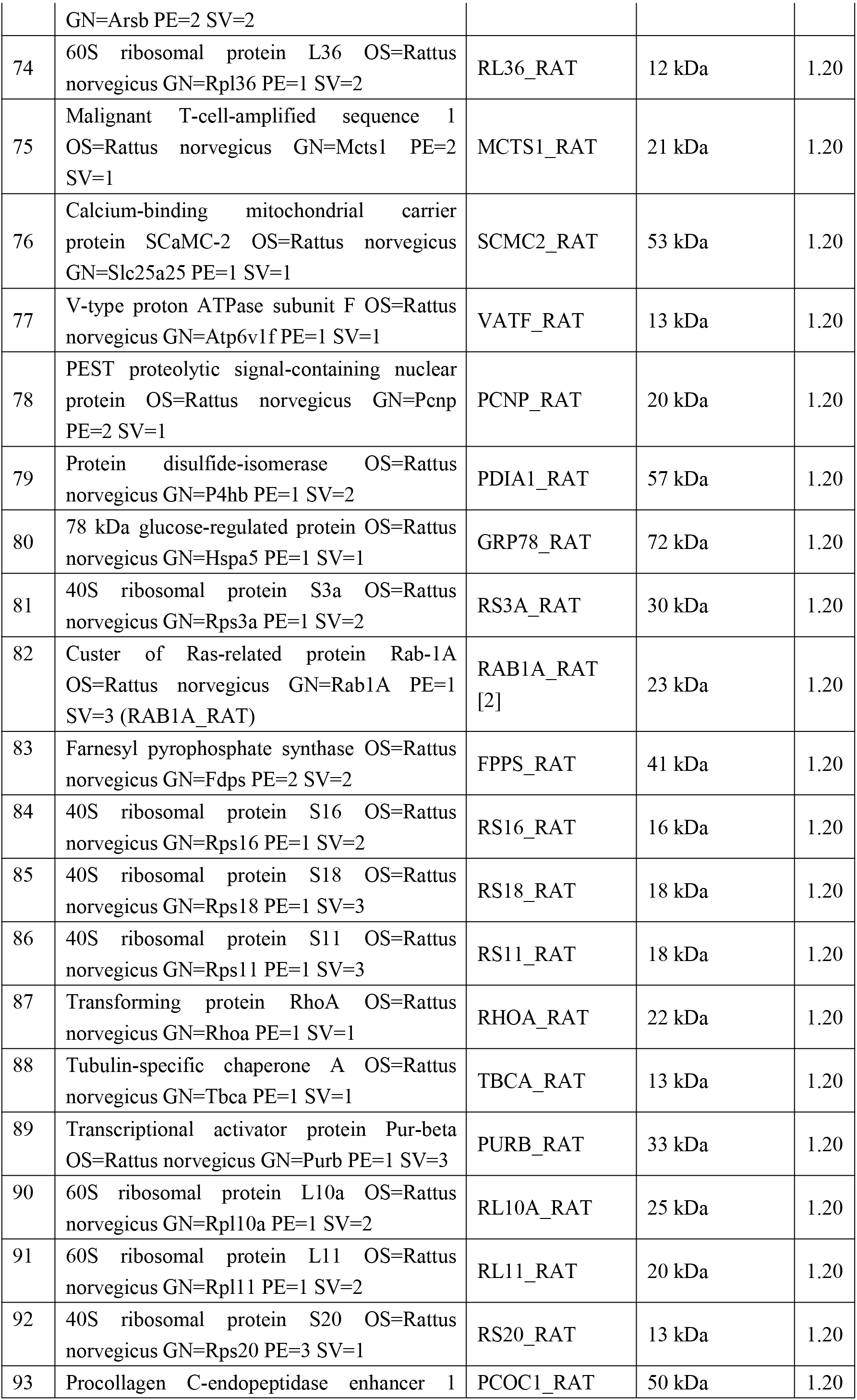
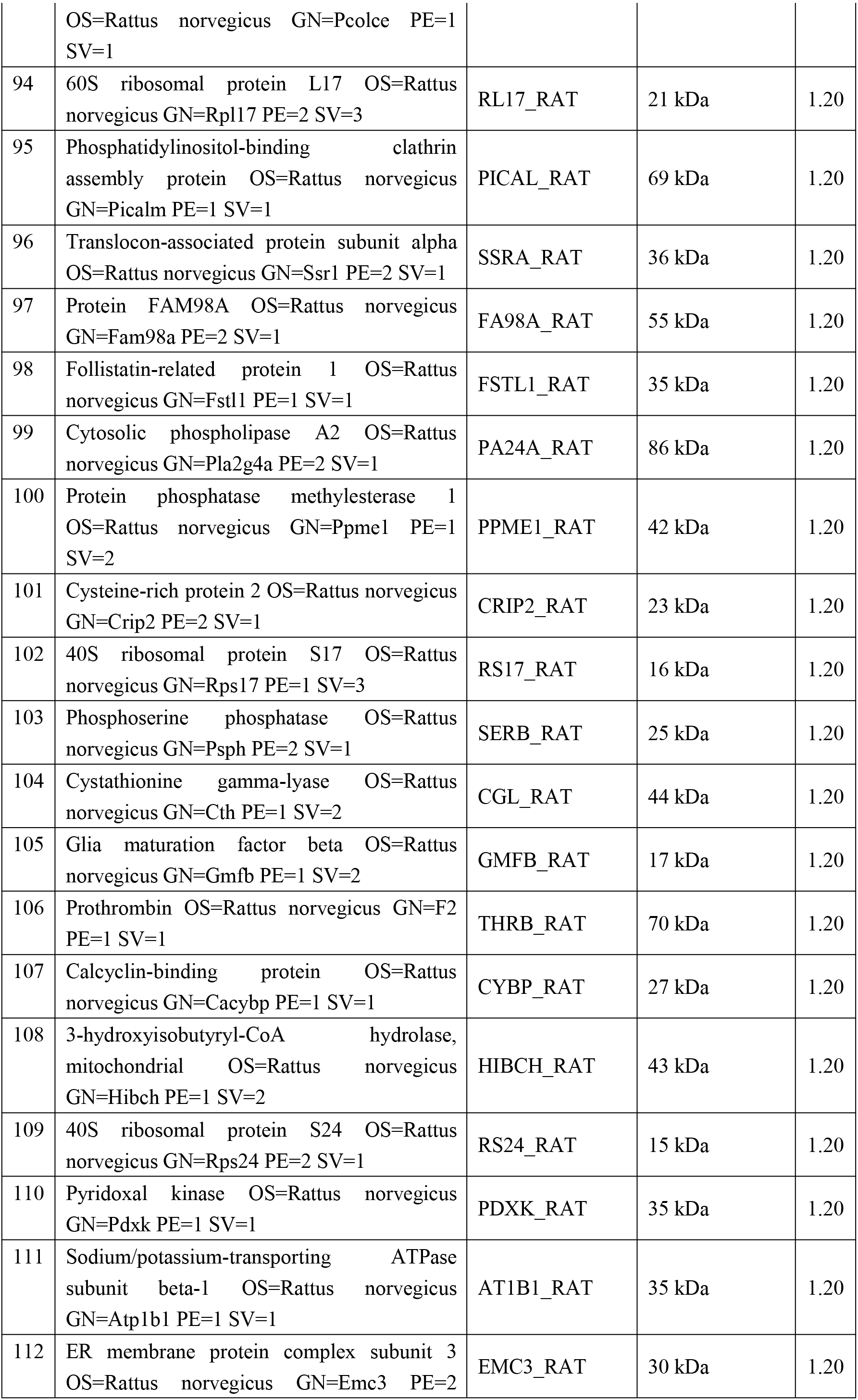
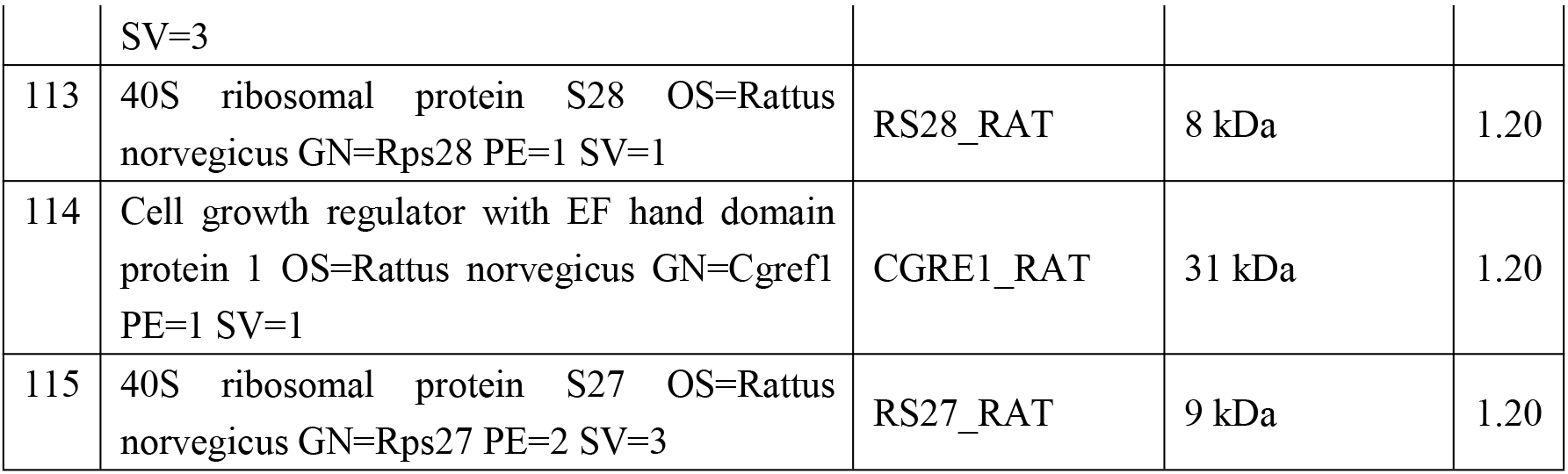
The up-regulation of differential proteins in fibroblasts induced by TGF-β1.

**Table S3.** GO analysis of differential proteins performed in PANTHER and classified based on molecular function.

| **molecular function** |  | **Up-regulated proteins** |  | **Down-regulated proteins** |
| --- | --- | --- | --- | --- |
| catalytic activity<br>(GO:0003824) | 37 | PDCL3; LYOX; TANC1; GMFB; CGL; FPPS; VATF; PDIA1; PCOC1; PPME1; ARHG6; LYPA2; SERB; TPM2; PDXK; ARSB; ERG1; ORN; EI2BG; RHOA; HIBCH; CK5P3; 2AAB; METK2; GFPT2; DYR; THRB; MAGI3; PAI1; PA24A; CYBP; ARFG1; ADAS; ATIF1; IPP2; GSH0; GNL3 | 28 | KIF1B; SMC1A; THTM; MRCKA; CAV1; BST1; CATL1; AL3A1; CATA; RPN2; CN37; CTHR1; LPP3; DHI1; PSME2; RASA1; COX7C; LPP1; P85A; GLO2; MAGT1; AAK1; BZW2; APOE; MYO1B; HXK1; NT5D2; HMGCL |
| binding<br>(GO:0005488) | 24 | GMFB; FGF2; CRIP2; SSRA; PCOC1; PAIRB; ARHG6; RL10A; S10AB; SERB; PURB; EI2BG; LRRF1; RHOA; FSTL1; CYR61; OLR1; TCTP; ARFG1; CHSP1; RS13; ATIF1; RL35A; YKT6 | 18 | SMC1A; CAV1; CATA; ADDG; MARE2; DTNB; H2AY; SQSTM; P85A; SPTN2; IBP5; AAK1; ADDA; C2D1A; APOE; H4; MYO1B; WNT5A |
| structural molecule activity<br>(GO:0005198) | 19 | RL11; RS11; CRIP2; RS20; RS12; RS28; RS16; RL10A; RS27; SCMC2; TPM2; RS18; RL23A; RL36; RL26; RS19; TCTP; RS13; RL35A | 14 | CAV1; M2OM; K2C1; ADDG; MGP; COL12; MARE2; DTNB; K2C4; MCAT; SPTN2; ADDA; MYO1B; MPCP |
| transporter activity<br>(GO:0005215) | 4 | CGRE1; VATF; PCOC1; AT1B1 | 7 | VDAC2; PLS3; SFXN3; VDAC3; ABCD3; APOE; CXA1 |
| receptor activity<br>(GO:0004872) | 3 | LYOX; PCOC1; MCTS1 | 4 | NTRI; ANTR1; MEP50; TPBG |
| translation regulator activity<br>(GO:0045182) | 1 | EI2BG | 2 | PDCD4; BZW2 |
| antioxidant activity | 0 |  | 1 | CATA |

**Table S4.**
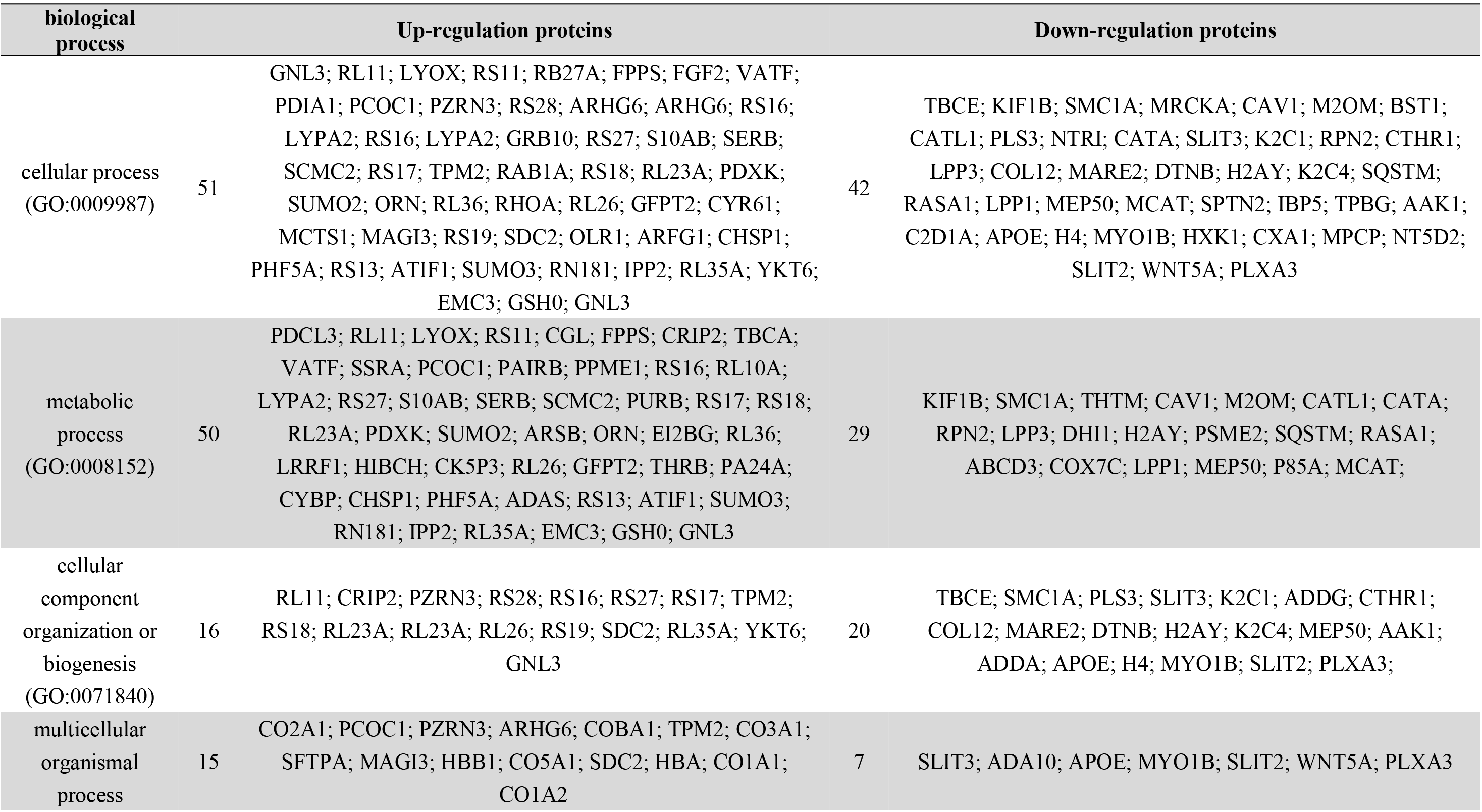
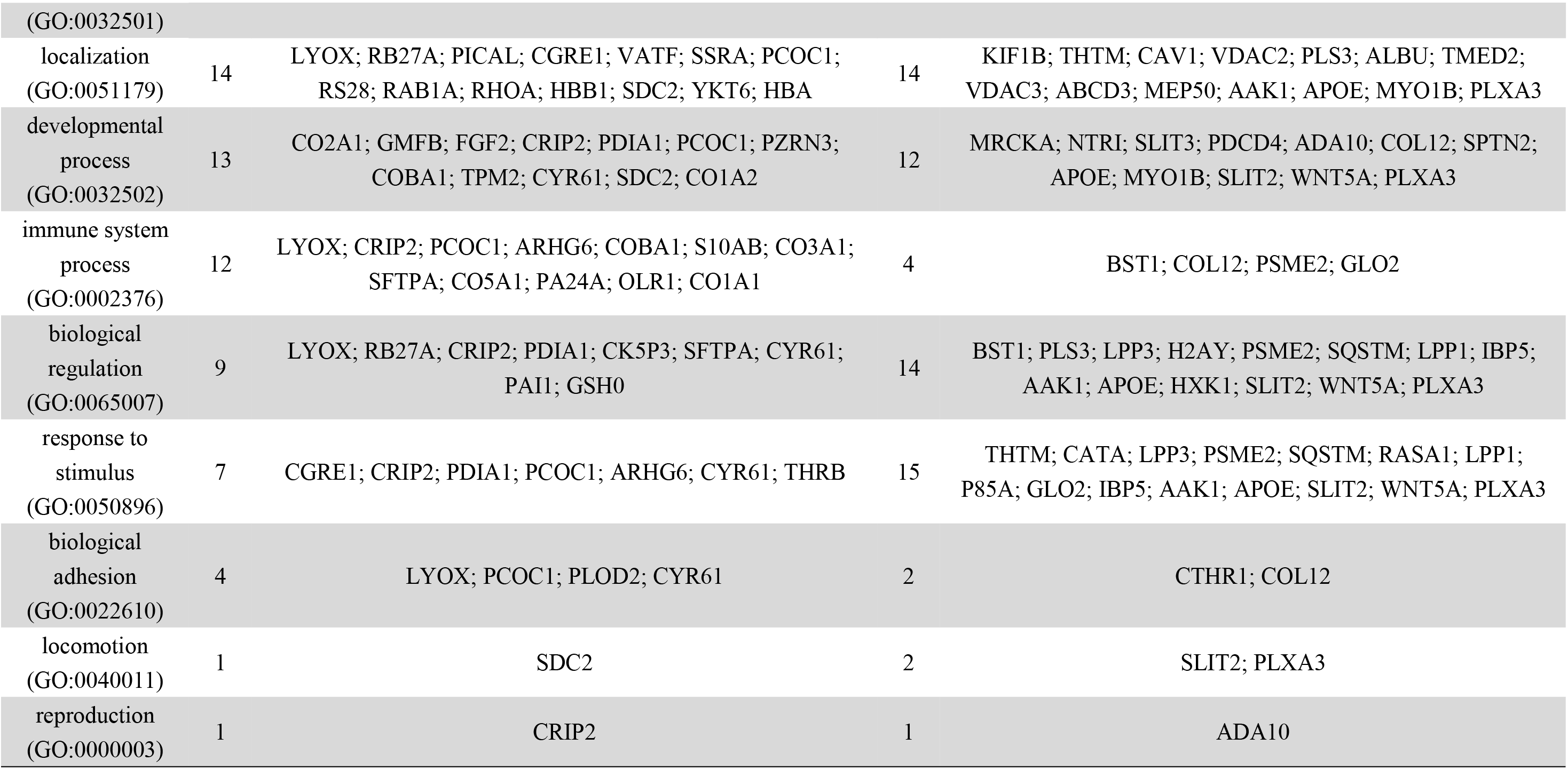
GO analysis of differential proteins performed in PANTHER and classified based on biological process.

**Table S5.** GO analysis of differential proteins performed in PANTHER and classified based on cellular component.

| **cellular component** |  | **Up-regulation proteins** |  | **Down-regulation proteins** |
| --- | --- | --- | --- | --- |
| cell part<br>(GO:0044464) | 43 | PDCL3; RL11; RS11; RB27A; PICAL; FPPS; CRIP2; RS20; RS12; NOLC1; RS28; RS16; FA98A; FA98A; LYPA2; RS27; SERB; RS17; TPM2; RAB1A; RS18; RL36A; RL23A; PDXK; SUMO2; RL36; CK5P3; RL26; METK2; RS19; SDC2; PA24A; TCTP; PHF5A; RS13; ATIF1; SUMO3; IPP2; RL35A; YKT6; EMC3; GSH0; RS3A; GNL3 | 23 | KIF1B; CATL1; CATA; K2C1; RPN2; RM37; ADDG; CN37; MARE2; DTNB; TMED2; H2AY; K2C4; PSME2; SQSTM; RASA1; P85A; SPTN2; AAK1; ADDA; C2D1A; MYO1B; HXK1 |
| organelle<br>(GO:0043226) | 26 | RL11; RS11; RB27A; RS20; RS12; NOLC1; RS28; RS16; RS27; RS17; RAB1A; RS18; RL36A; RL23A; SUMO2; RL36; RL26; RS19; TCTP; PHF5A; RS13; ATIF1; SUMO3; RL35A; EMC3; GNL3 | 15 | KIF1B; CATL1; K2C1; RPN2; RM37; ADDG; MARE2; DTNB; H2AY; K2C4; PSME2; SQSTM; ADDA; C2D1A; MYO1B |
| macromolecular complex<br>(GO:0032991) | 24 | RL11; RS11; PICAL; RS20; RS12; RS28; RS16; FA98A; RS27; RS17; RS18; RL36A; RL23A; RL36; RL26; METK2; RS19; PHF5A; RS13; IPP2; RL35A; YKT6; EMC3; GSH0 | 6 | KIF1B; RPN2; TMED2; PSME2; P85A; APOE |
| membrane<br>(GO:0016020) | 4 | RAB1A; MAGI3; YKT6; EMC3 | 8 | BST1; PLS3; RPN2; LPP3; RASA1; LPP1; MYO1B; CXA1 |
| extracellular region<br>(GO:0005576) | 3 | PCOC1; CYR61; PAI1 | 6 | CATL1; COL12; IBP5; TPBG; APOE; WNT5A |
| extracellular matrix<br>(GO:0031012) | 2 | PCOC1; CYR61 |  | MMP2; CTHR1; COL12 |
| cell junction<br>(GO:0030054) | 1 | MAGI3 | 1 | MYO1B |
| synapse<br>(GO:0045202) | 1 | PZRN3 | 0 |  |

**Table S6.**
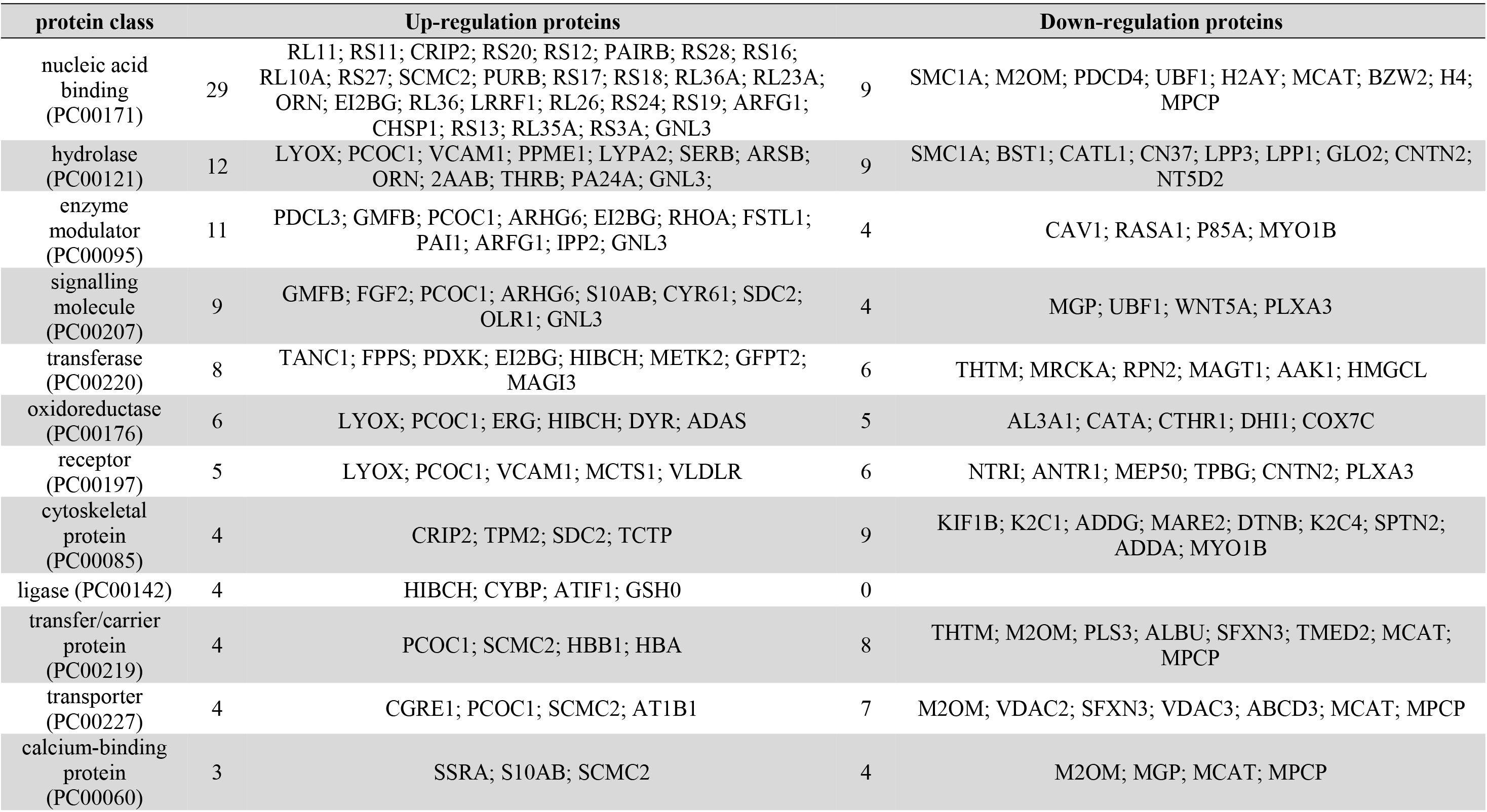
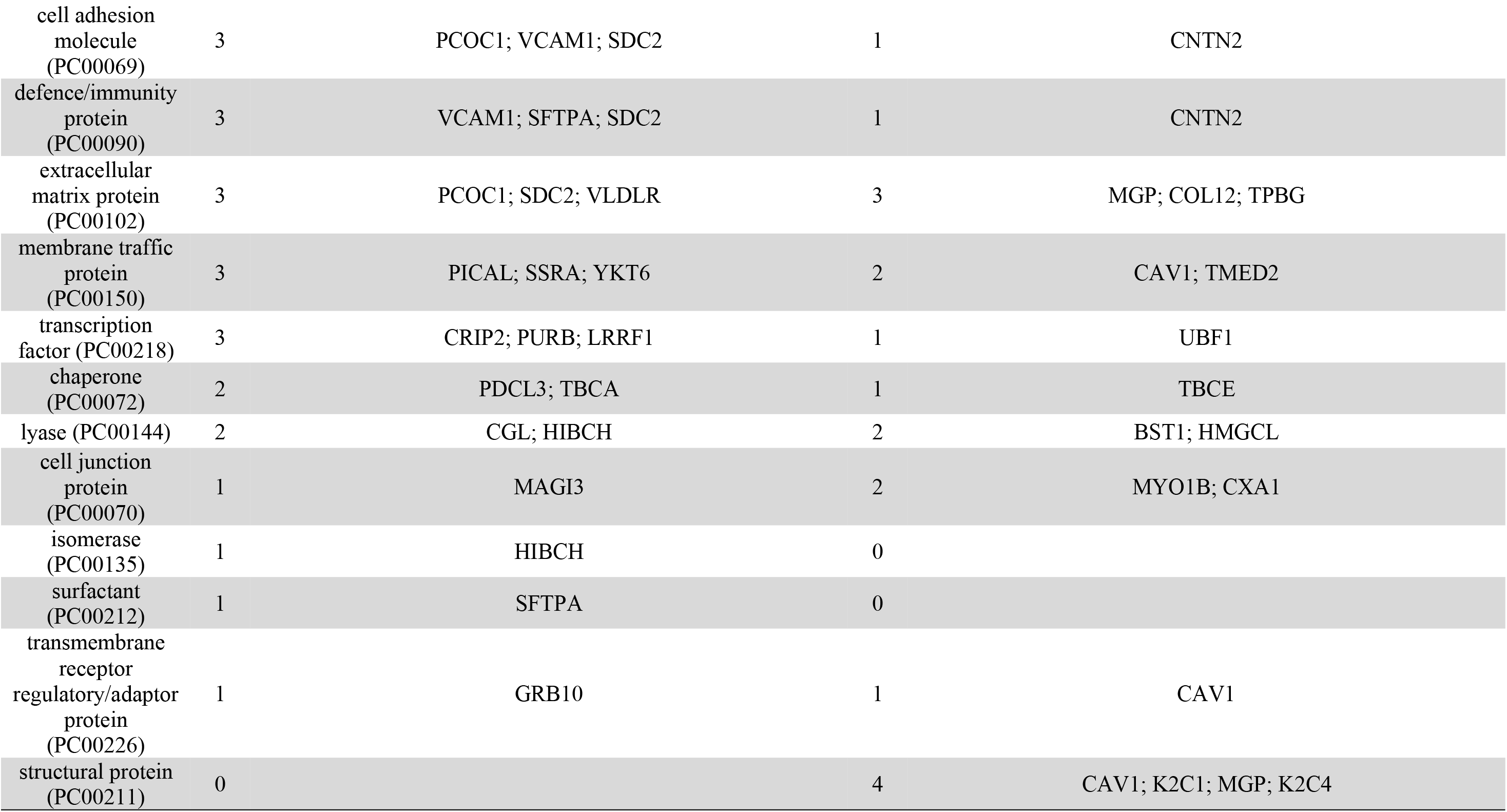
Protein class ontology analysis of differential proteins performed in PANTHER.

| protein class |  | Up-regulation proteins |  | Down-regulation proteins |
| --- | --- | --- | --- | --- |
| nucleic acid binding (PC00171) | 29 | RL11; RS11; CRIP2; RS20; RS12; PAIRB; RS28; RS16; RL10A; RS27; SCMC2; PURB; RS17; RS18; RL36A; RL23A; ORN; EI2BG; RL36; LRRF1; RL26; RS24; RS19; ARFG1; CHSP1; RS13; RL35A; RS3A; GNL3 | 9 | SMC1A; M2OM; PDCD4; UBF1; H2AY; MCAT; BZW2; H4; MPCP |
| hydrolase (PC00121) | 12 | LYOX; PCOC1; VCAM1; PPME1; LYPA2; SERB; ARSB; ORN; 2AAB; THRB; PA24A; GNL3; | 9 | SMC1A; BST1; CATL1; CN37; LPP3; LPP1; GLO2; CNTN2; NT5D2 |
| enzyme modulator (PC00095) | 11 | PDCL3; GMFB; PCOC1; ARHG6; EI2BG; RHOA; FSTL1; PAI1; ARFG1; IPP2; GNL3 | 4 | CAV1; RASA1; P85A; MYO1B |
| signalling molecule (PC00207) | 9 | GMFB; FGF2; PCOC1; ARHG6; S10AB; CYR61; SDC2; OLR1; GNL3 | 4 | MGP; UBF1; WNT5A; PLXA3 |
| transferase (PC00220) | 8 | TANC1; FPPS; PDXK; EI2BG; HIBCH; METK2; GFPT2; MAGI3 | 6 | THTM; MRCKA; RPN2; MAGT1; AAK1; HMGCL |
| oxidoreductase (PC00176) | 6 | LYOX; PCOC1; ERG; HIBCH; DYR; ADAS | 5 | AL3A1; CATA; CTHR1; DHI1; COX7C |
| receptor (PC00197) | 5 | LYOX; PCOC1; VCAM1; MCTS1; VLDLR | 6 | NTRI; ANTR1; MEP50; TPBG; CNTN2; PLXA3 |
| cytoskeletal protein (PC00085) | 4 | CRIP2; TPM2; SDC2; TCTP | 9 | KIF1B; K2C1; ADDG; MARE2; DTNB; K2C4; SPTN2; ADDA; MYO1B |
| ligase (PC00142) | 4 | HIBCH; CYBP; ATIF1; GSH0 | 0 |  |
| transfer/carrier protein (PC00219) | 4 | PCOC1; SCMC2; HBB1; HBA | 8 | THTM; M2OM; PLS3; ALBU; SFXN3; TMED2; MCAT; MPCP |
| transporter (PC00227) | 4 | CGRE1; PCOC1; SCMC2; AT1B1 | 7 | M2OM; VDAC2; SFXN3; VDAC3; ABCD3; MCAT; MPCP |
| calcium-binding protein (PC00060) | 3 | SSRA; S10AB; SCMC2 | 4 | M2OM; MGP; MCAT; MPCP |
| cell adhesion molecule (PC00069) | 3 | PCOC1; VCAM1; SDC2 | 1 | CNTN2 |
| defence/immunity protein (PC00090) | 3 | VCAM1; SFTPA; SDC2 | 1 | CNTN2 |
| extracellular matrix protein (PC00102) | 3 | PCOC1; SDC2; VLDLR | 3 | MGP; COL12; TPBG |
| membrane traffic protein (PC00150) | 3 | PICAL; SSRA; YKT6 | 2 | CAV1; TMED2 |
| transcription factor (PC00218) | 3 | CRIP2; PURB; LRRF1 | 1 | UBF1 |
| chaperone (PC00072) | 2 | PDCL3; TBCA | 1 | TBCE |
| lyase (PC00144) | 2 | CGL; HIBCH | 2 | BST1; HMGCL |
| cell junction protein (PC00070) | 1 | MAGI3 | 2 | MYO1B; CXA1 |
| isomerase (PC00135) | 1 | HIBCH | 0 |  |
| surfactant (PC00212) | 1 | SFTPA | 0 |  |
| transmembrane receptor regulatory/adaptor protein (PC00226) | 1 | GRB10 | 1 | CAV1 |
| structural protein (PC00211) | 0 |  | 4 | CAV1; K2C1; MGP; K2C4 |

**Table S7.**
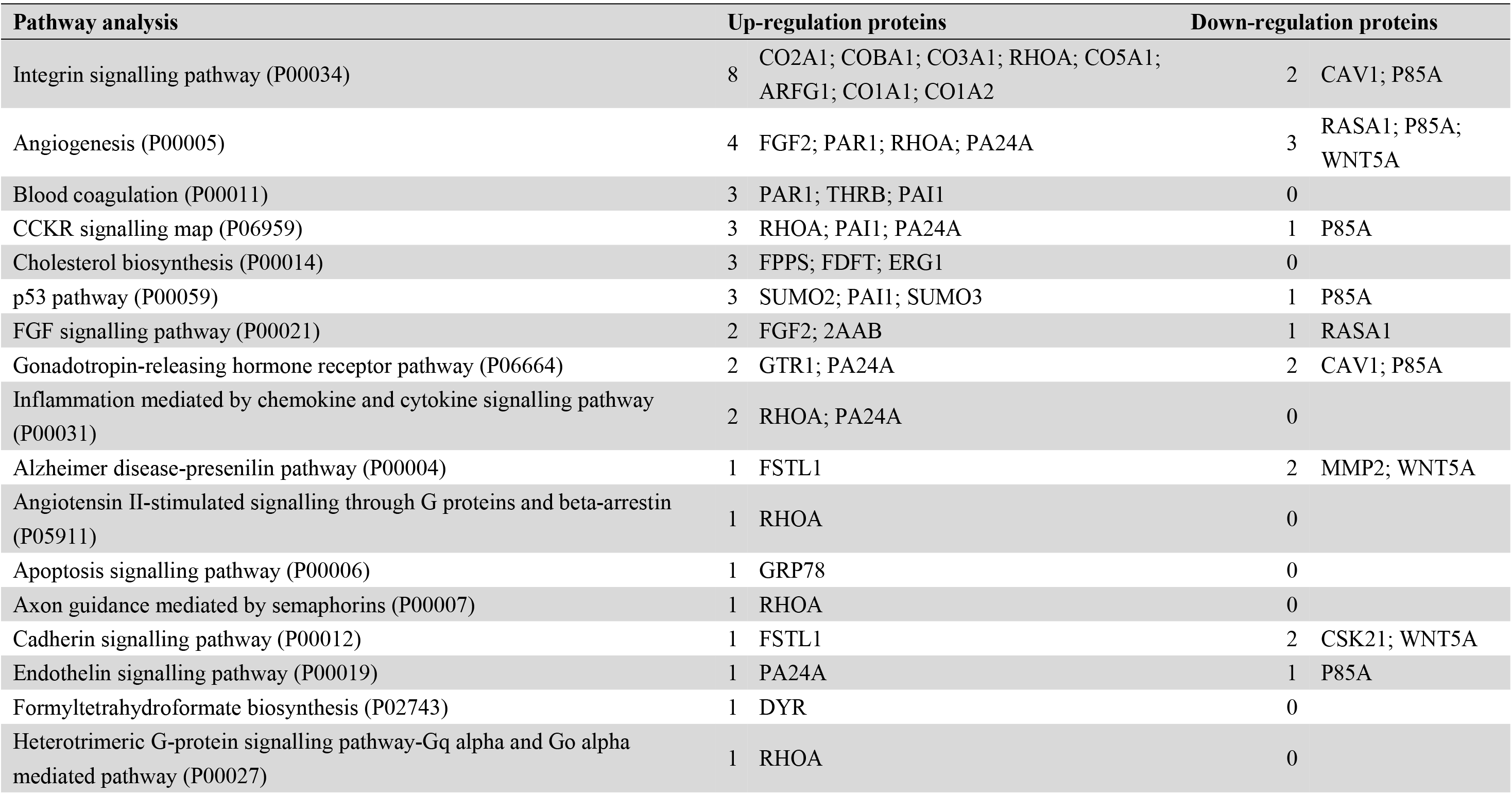
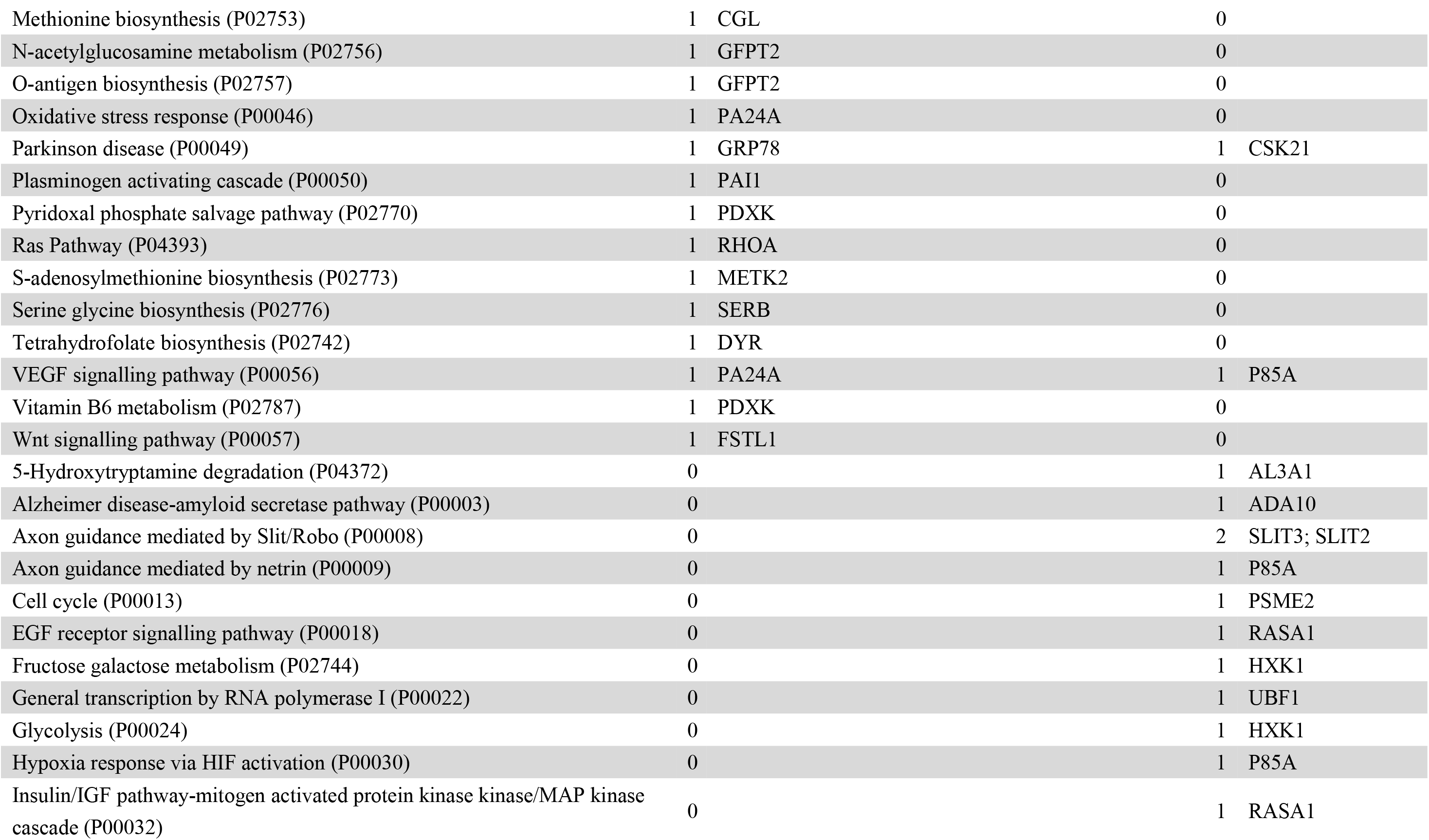
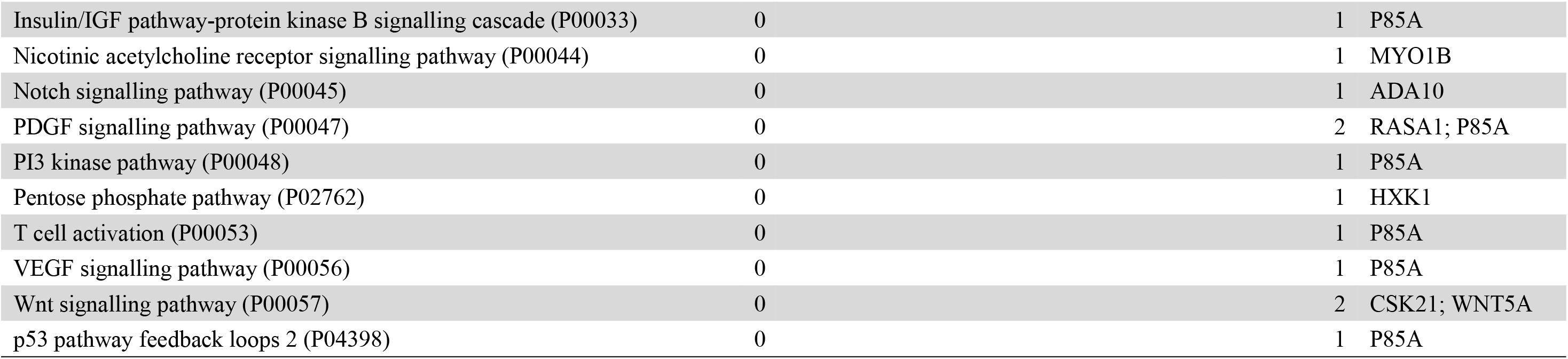
Pathway analysis of differential proteins performed in PANTHER.

